# Deconvolution of polygenic risk score in single cells unravels cellular and molecular heterogeneity of complex human diseases

**DOI:** 10.1101/2024.05.14.594252

**Authors:** Sai Zhang, Hantao Shu, Jingtian Zhou, Jasper Rubin-Sigler, Xiaoyu Yang, Yuxi Liu, Johnathan Cooper-Knock, Emma Monte, Chenchen Zhu, Sharon Tu, Han Li, Mingming Tong, Joseph R. Ecker, Justin K. Ichida, Yin Shen, Jianyang Zeng, Philip S. Tsao, Michael P. Snyder

## Abstract

Polygenic risk scores (PRSs) are commonly used for predicting an individual’s genetic risk of complex diseases. Yet, their implication for disease pathogenesis remains largely limited. Here, we introduce scPRS, a geometric deep learning model that constructs single-cell-resolved PRS leveraging reference single-cell chromatin accessibility profiling data to enhance biological discovery as well as disease prediction. Real-world applications across multiple complex diseases, including type 2 diabetes (T2D), hypertrophic cardiomyopathy (HCM), and Alzheimer’s disease (AD), showcase the superior prediction power of scPRS compared to traditional PRS methods. Importantly, scPRS not only predicts disease risk but also uncovers disease-relevant cells, such as hormone-high alpha and beta cells for T2D, cardiomyocytes and pericytes for HCM, and astrocytes, microglia and oligodendrocyte progenitor cells for AD. Facilitated by a layered multi-omic analysis, scPRS further identifies cell-type-specific genetic underpinnings, linking disease-associated genetic variants to gene regulation within corresponding cell types. We substantiate the disease relevance of scPRS-prioritized HCM genes and demonstrate that the suppression of these genes in HCM cardiomyocytes is rescued by Mavacamten treatment. Additionally, we establish a novel microglia-specific regulatory relationship between the AD risk variant rs7922621 and its target genes *ANXA11* and *TSPAN14*. We further illustrate the detrimental effects of suppressing these two genes on microglia phagocytosis. Our work provides a multi-tasking, interpretable framework for precise disease prediction and systematic investigation of the genetic, cellular, and molecular basis of complex diseases, laying the methodological foundation for single-cell genetics.

## Main

Polygenic risk score (PRS)^1^, also known as polygenic score (PGS)^2^, is a standard approach for predicting quantitative traits and disease risks based on an individual’s genetic makeup. The method is built upon genetic variants including single nucleotide polymorphisms (SNPs) and small insertions and deletions (INDELs) which are common (minor allele frequency (MAF) > 5%) in the population. PRS is a critical component of precision genomic medicine and has promised versatile utilities^3^, such as health management, disease screening, and therapeutic intervention. Traditionally, PRS computation involves a linear model that sums the genotypes of selected variants, with each variant weighted according to its effect size as determined by the genome-wide association study (GWAS)^4^. The clumping and thresholding (C+T) method serves as the basis of constructing PRSs, but various advanced approaches^5–8^ have been developed to enhance prediction by considering nuanced genetic architectures.

Complex diseases exhibit significant cellular heterogeneity, involving multiple tissues or cell types in their pathogenesis^9^. Risk variants, particularly noncoding ones, can influence disease susceptibility and phenotypic variability through diverse cellular and molecular processes^10–12^. However, these multiple layers of complexity have been oversimplified in conventional modeling, substantially limiting the prediction power and interpretability of PRS^13^. On the other hand, single-cell sequencing has emerged as a potent tool to dissect cellular and molecular heterogeneity across different tissues and conditions^14^, offering unprecedented opportunities to explore genome function at high resolution.

To address these challenges, here we proposed a novel strategy that unifies genetics and single-cell genomics, named single-cell genetics^15^, for studying disease genetics at single-cell resolution. In particular, we introduced **<u>s</u>**ingle-**<u>c</u>**ell **<u>p</u>**olygenic **<u>r</u>**isk **<u>s</u>**core (scPRS), a geometric deep learning model^16^ that enables individualized genetic risk prediction at the single-cell level. scPRS leverages a graph neural network (GNN) to construct genetic risk score by drawing insights from reference single-cell chromatin accessibility measured by single-cell or single-nuclei sequencing assay for transposase-accessible chromatin (scATAC-seq or snATAC-seq)^17^. Beyond enhanced disease prediction, scPRS is empowered with fine-grained model interpretability, which allows for systematic discovery of cell types and cell-type-specific gene regulatory programs underpinning diseases.

We performed extensive simulations demonstrating the effectiveness and robustness of scPRS in identifying phenotype-relevant cells. We applied scPRS to three complex diseases – type 2 diabetes (T2D), hypertrophic cardiomyopathy (HCM), and Alzheimer’s disease (AD) – and showcased its superior prediction performance compared to multiple traditional PRS methods. Using model interpretation, scPRS identified known disease-critical cell types and also discovered novel cell populations relevant to each disease. For instance, the cells whose roles in specific diseases were less well recognized, including alpha cells, pericytes, and oligodendrocyte progenitor cells (OPCs), were strongly linked to T2D, HCM, and AD, respectively. scPRS-powered functional analysis further pinpointed candidate causal variants and mapped them to cis-regulatory elements (CREs) and target genes within specific cell types, unraveling a cell-type-specific landscape of genetic regulation. Using drug perturbation data, we validated our scPRS-prioritized HCM genes, showing that the suppression of these genes in diseased cardiomyocytes was rescued by Mavacamten (an FDA-approved HCM drug) treatment. Supported by experiments, we illustrated a new role of the AD risk variant rs7922621 in down-regulating two downstream genes, *ANXA11* and *TSPAN14*, specifically in microglia, and further revealed the phenotypic effects of suppression of these two genes on impairing microglia phagocytosis.

## Results

### Overview of scPRS

Our design principle of scPRS is to leverage single-cell epigenome profiling to rationalize the calculation of polygenic risk scores. The approach begins with deconvoluting traditional PRS within individual cells based on their chromatin accessibility profiled by single-cell ATAC-seq, followed by the integration of deconvoluted cell-level PRSs into a final score capitalizing on cell-cell similarities (**Fig. 1**; **Methods**). In particular, utilizing GWAS summary statistics derived from a disease cohort (referred to as the discovery cohort) and a single-cell ATAC-seq dataset of healthy tissue (referred to as the reference dataset) pertinent to the disease, we compute a conditioned PRS for each individual within our target cohort (not overlapping the discovery cohort) and for each reference cell, in which we mask out genetic variants located outside open chromatin regions captured in that specific cell. Recognizing the sparsity of single-cell ATAC-seq signals, scPRS further refines per-cell PRS features by employing a graph neural network (GNN)^18^. This GNN operation serves the dual purpose of denoising raw PRS profiles and capturing non-linear relationships between genetic signals and cellular epigenome. This extends the linear structure adopted by traditional PRS methods. In the final step, scPRS aggregates smoothed cell-level PRSs and yields a final score that indicates disease risk. The interpretability of scPRS is achieved by learned model weights accompanied with individual cells (**Fig. 1**; **Methods**) which indicate the contribution of different cells to disease risk.

**Fig. 1.**
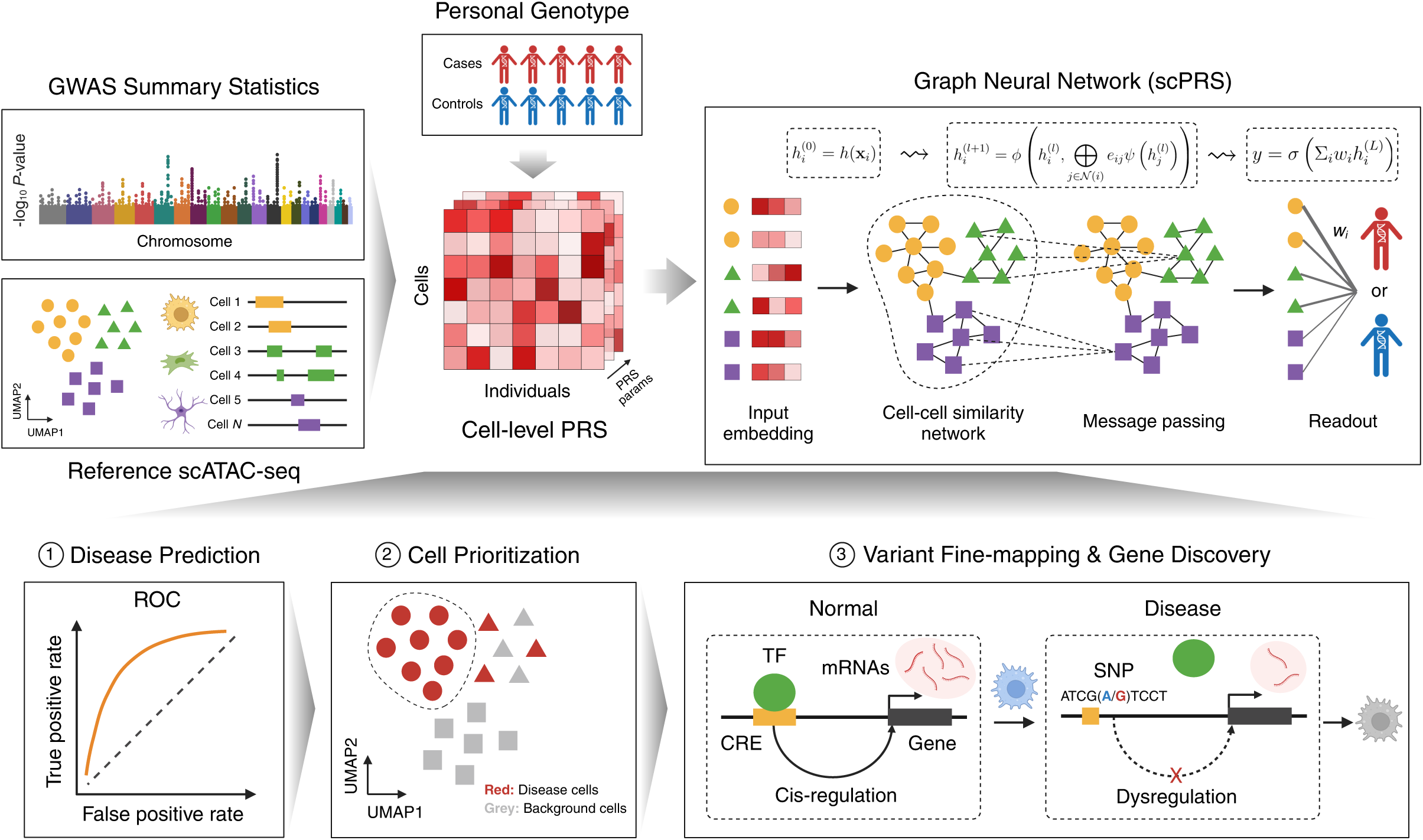
Overview of the scPRS framework and its applications. For a given complex disease, scPRS first leverages GWAS summary statistics derived from the discovery cohort and the reference single-cell ATAC-seq dataset to calculate cell-level PRSs with different PLINK parameters for individuals in the target cohort. Next, scPRS embeds and propagates cell-level PRSs over the cell-cell similarity network using a graph neural network (GNN). The final readout combines smoothed PRSs from all cells to predict the disease status. scPRS is trained to minimize the error between predicted and true disease status. The trained model can be used to (1) predict disease risk for unseen individuals, (2) prioritize disease-relevant cells and cell types, and (3) fine-map disease risk variants, pinpoint candidate disease genes, and reveal disrupted gene regulation in specific cell types. GWAS, genome-wide association study; UMAP, uniform manifold approximation and projection; params, parameters; ROC, receiver operating characteristic; TF, transcription factor; CRE, cis-regulatory element; SNP, single nucleotide polymorphism; mRNAs, messenger RNAs.

The functionality of scPRS is exemplified by three downstream tasks. First, scPRS accurately predicts disease risk for unseen individuals solely based on their genotypes (**Fig. 1**, Panel 1). Second, scPRS systematically prioritizes single-cells that are relevant to the disease, overcoming the resolution constraint of cell clustering (**Fig. 1**, Panel 2). Third, facilitated by a multiomic approach, scPRS fine-maps causal variants, pinpoints candidate genes, and identifies disease-relevant gene dysregulation within prioritized cell types (**Fig. 1**, Panel 3). These applications highlight the broad utilities of scPRS in dissecting genetic, cellular, and molecular heterogeneity of complex diseases.

### Evaluation of scPRS using simulations

We first performed simulation experiments to evaluate the capacity of scPRS in identifying phenotype-relevant cells. Assuming that the trait “monocyte count” is fully determined by genetic variants located within monocyte-specific open chromatin^19^, we simulated monocyte counts for individuals of a genotyped cohort^20^ (*n* = 401; **Methods**). We then asked whether we could use scPRS to recapitulate monocytes as the causal cell type. Specifically, we used a reference scATAC-seq dataset^21^ (**Fig. 2a**) of human peripheral blood mononuclear cells (PBMCs) to identify monocyte-specific peaks (**Methods**). Based on a monocyte count GWAS^19^ defining variant effect sizes, we simulated monocyte count for each individual by calculating the clumping and thresholding (C+T) PRS using only variants within monocyte-specific peaks (**Methods**). Next, we trained an scPRS model to predict simulated monocyte counts from cell-level PRSs computed on all PBMCs. We observed that scPRS predictions were significantly correlated with simulated monocyte counts (*r* = 0.77, *P* < 2.2 × 10^−16^, Pearson correlation; **Fig. 2b**). Notably, the cells prioritized by our model (**Methods**) were significantly enriched within monocytes (*Z* = 39.58, *P* < 1 × 10^−50^, two-sided Fisher’s exact test; **Fig. 2c**), indicating that scPRS captured causal cells.

**Fig. 2.**
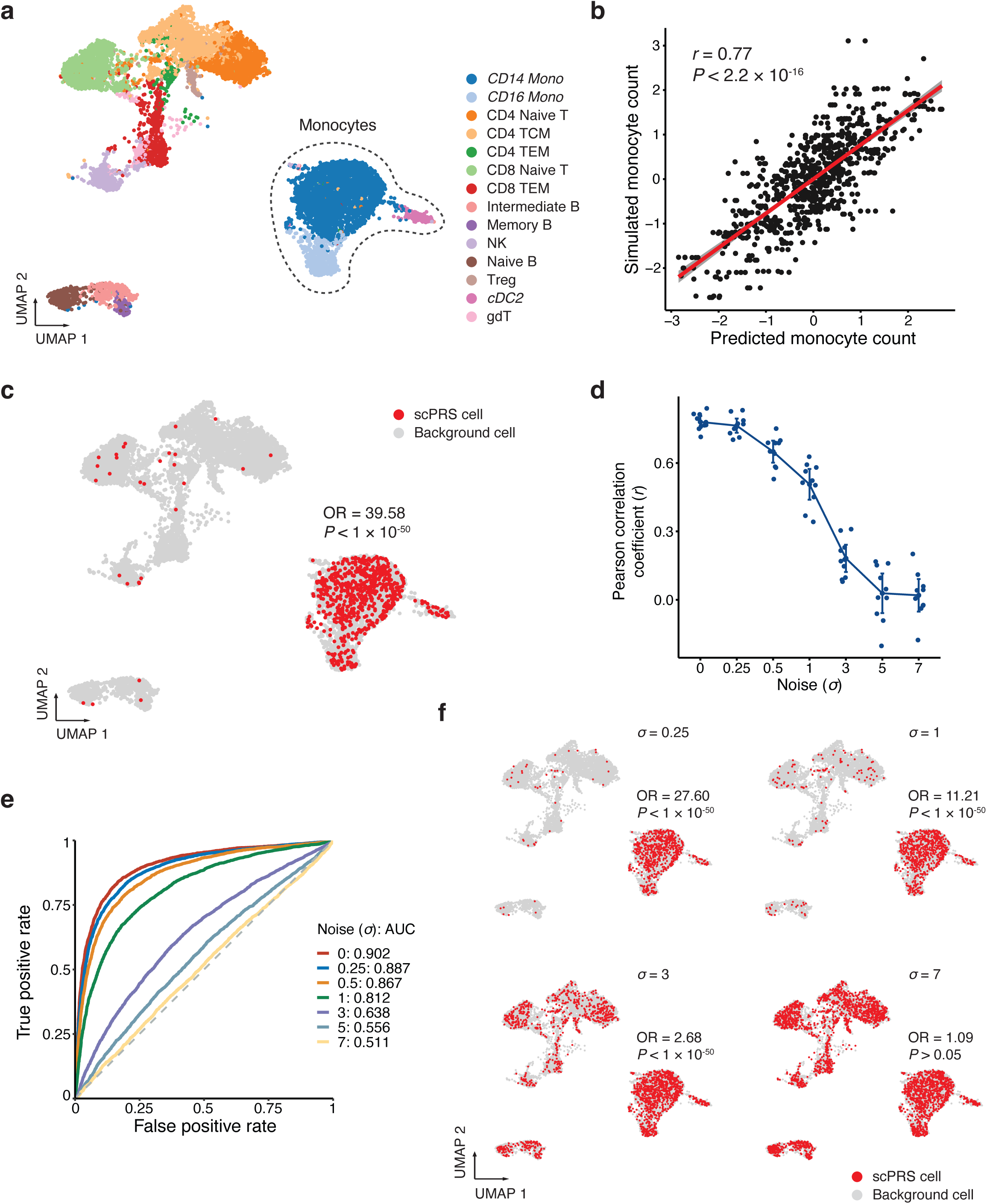
Assessing performance of scPRS using simulations. **a**, The uniform manifold approximation and projection (UMAP) plot of the human peripheral blood mononuclear cell (PBMC) scATAC-seq dataset. Cell clusters with less than 150 cells are not shown. Monocyte subtypes are highlighted in italic. Mono, monocyte; TCM, memory T cell; TEM, effector memory T cell; NK, natural killer cell; Treg, regulatory T cell; cDC2, conventional type 2 dendritic cell; gdT, gamma-delta T cell. **b**, Pearson correlation between simulated and predicted monocyte counts (*n* = 10 repeats). The linear regression line and 95% confidence interval (CI) are annotated in the red line and gray shaded area, respectively. **c**, Monocyte-count-relevant cells prioritized by scPRS (in red). Odds ratio and *P*-value by two-sided Fisher’s exact test. OR, odds ratio. **d**, Pearson correlation between simulated and predicted monocyte counts (*n* = 10 repeats) in different noise settings. The mean and 95% CI are annotated in the dot and error bar, respectively. **e**, The receiver operating characteristic (ROC) curves for cell prioritization in different noise settings, wherein monocytes were labeled as “1” and other cells were labeled as “0”. AUC, the area under the curve. **f**, Monocyte-count-relevant cells prioritized by scPRS (in red) in different noise settings. Odds ratio and *P*-value by two-sided Fisher’s exact test.

Human phenotypes such as complex diseases can be influenced by various non-genetic factors, including environmental and lifestyle factors^22^. Additionally, the measurement of phenotypes often carries inherent noise. Therefore, it is important to assess the robustness of scPRS by introducing noise and randomness into the simulation (**Methods**). As expected, we observed a progressive reduction in prediction performance as we introduced larger amounts of noise (**Fig. 2d**). Surprisingly, scPRS sustained its ability to uncover monocytes even in the presence of considerable noise terms (**Fig. 2e,f** and **Extended Data Fig. 1**). For example, when we introduced a noise term with the same amount of randomness (standard deviation (SD) *σ* = 1) as that of the simulated phenotype, scPRS still accurately identified monocytes (area under the curve (AUC) = 0.812; **Fig. 2e**); the enrichment of monocytes persisted even when three times the amount of randomness was added (*σ* = 3; *Z* = 2.68, *P* < 1 × 10^−50^, two-sided Fisher’s exact test; **Fig. 2f**). These results demonstrated the efficacy and robustness of scPRS in identifying phenotype-relevant cell types, even in the context of multifactorial phenotypes or noisy data.

### scPRS accurately predicts diseases

We applied scPRS to several diseases, including type 2 diabetes (T2D), hypertrophic cardiomyopathy (HCM), and Alzheimer’s disease (AD). We used UK Biobank (UKBB)^23^ data to construct target cohorts for T2D and AD (**Methods**), and our in-house whole-genome sequencing (WGS) data^24^ for HCM (**Methods**). The discovery GWAS dataset was carefully chosen to ensure non-overlap with the target cohort for each disease^25–27^. Multiple reference single-cell ATAC-seq datasets of disease-relevant tissues were used, including pancreas^28^ for T2D, left ventricle^29^ for HCM, and frontal cortex^30^ for AD (**Methods**).

For benchmarking, we employed five well-established PRS methods, including C+T (implemented by PLINK^31^), LDpred2 (LDpred2-inf, LDpred2-grid, and LDpred2-auto)^5^, and Lassosum^7^ (**Methods**). To examine the predictability of non-peak and non-genetic factors, we also built a C+T PRS model based on variants situated beyond open chromatin regions, and a logistic regression model using age, sex, and first 10 principal components (PCs) as input features (**Methods**).

Remarkably, scPRS-based methods consistently outperformed all other PRS approaches in all three diseases (**Fig. 3** and **Extended Data Fig. 2**). In particular, for HCM and AD, scPRS achieved significantly superior prediction performance based on both the area under the receiver operating characteristic curve (AUROC; HCM: mean = 0.692, SD = 0.079; AD: mean = 0.743, SD = 0.017) and the area under the precision-recall curve (AUPRC; HCM: mean = 0.781, SD = 0.062; AD: mean = 0.751, SD = 0.035), compared to all traditional PRS methods (adjusted *P* < 0.1, Benjamini-Hochberg (BH) correction; **Fig. 3b,c** and **Extended Data Fig. 2b**,c), except that scPRS exhibited comparable AUPRC to that of C+T for AD (*P* > 0.05, one-sided paired *t*-test; **Extended Data Fig. 2c**).

**Fig. 3.**
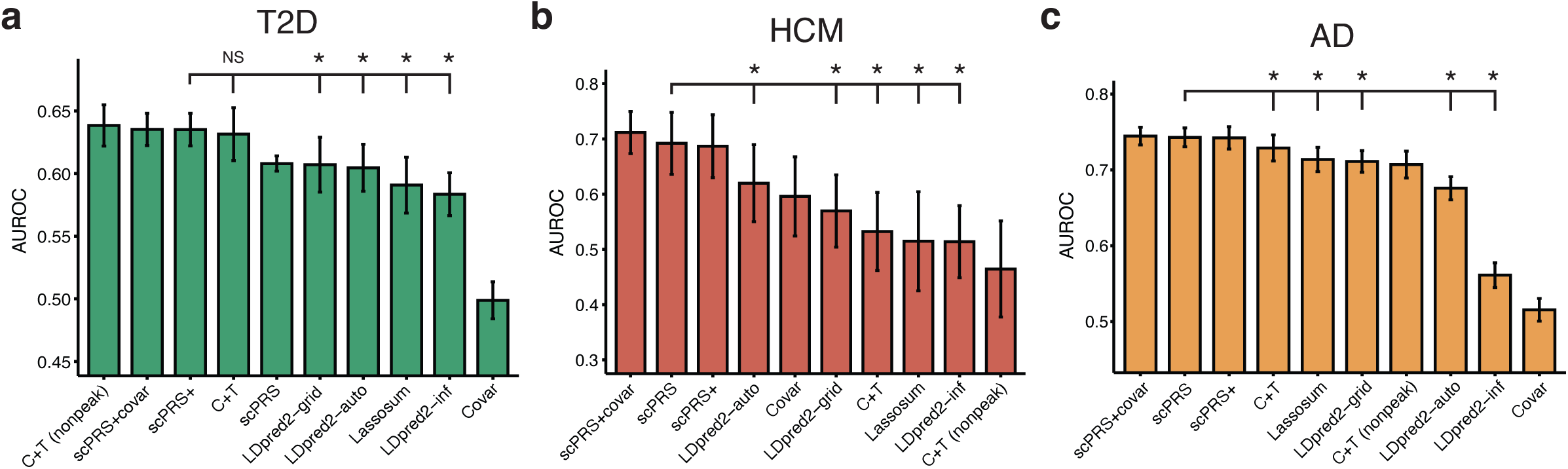
Prediction performance of scPRS and other baseline methods. **a-c**, Barplots of AUROC scores (*n* = 10 repeats) of different models for T2D (**a**), HCM (**b**), and AD (**c**), respectively. Training, validation, and test data splits were kept identical across different methods to ensure comparison fairness. scPRS+, scPRS model integrating non-peak PRSs; scPRS+covar, scPRS model integrating non-peak PRSs and covariates (i.e., age, sex, and 10 principle components); C+T, clumping and thresholding PRS; C+T (nonpeak), logistic regression model of non-peak C+T PRSs; Covar, logistic regression model of covariates. Performance comparison was conducted using one-sided paired *t*-test. AUROC, the area under the receiver operating characteristic curve; *, adjusted *P* < 0.1; NS, not significant. The mean and 95% confidence interval (CI) are annotated using the barplot and error bar, respectively.

For T2D, scPRS yielded performance comparable to other methods (AUROC: mean = 0.608, SD = 0.009; AUPRC: mean = 0.598, SD = 0.032; **Fig. 3a** and **Extended Data Fig. 2a**). Interestingly, after integrating non-peak C+T PRSs into the scPRS model (scPRS+; **Methods**), we obtained significantly superior prediction performance (AUROC: mean = 0.635, SD = 0.018; AUPRC: mean = 0.633, SD = 0.036), in comparison to all other PRS methods (adjusted *P* < 0.1, BH correction; **Fig. 3a** and **Extended Data Fig. 2a**), with the exception of C+T where the AUROC remained comparable (*P* > 0.05, one-sided paired *t*-test; **Fig. 3a**). These results suggest that the variants located outside pancreas open chromatin regions, such as protein-coding^32^ and splicing^33^ variants, or variants within CREs specific to other tissues^34^, may also contribute to T2D susceptibility. This is also supported by the observation that a predictor built solely on non-peak PRSs (C+T (nonpeak)) was ranked highest in both AUROC and AUPRC performance (AUROC: mean = 0.638, SD = 0.023; AUPRC: mean = 0.633, SD = 0.039; **Fig. 3a** and **Extended Data Fig. 2a**).

The covariate models exhibited limited prediction power for T2D (AUROC: mean = 0.499, SD = 0.021; AUPRC: mean = 0.502, SD = 0.038; **Fig. 3a** and **Extended Data Fig. 2a**) and AD (AUROC: mean = 0.515, SD = 0.021; AUPRC: mean = 0.524, SD = 0.043; **Fig. 3c** and **Extended Data Fig. 2c**) due to the fact that we matched age, sex, and population between cases and controls in constructing the target cohorts. Not surprisingly, the prediction performance was further boosted for all diseases after integrating all other factors, including non-peak PRSs and covariates, into the scPRS model (scPRS+covar; **Fig. 3** and **Extended Data Fig. 2**).

### scPRS identifies disease-relevant cells

Next, we sought to examine the disease-cell association using scPRS. For each disease, we first trained 100 scPRS models based on the entire cohort, and then we prioritized cells whose model weights consistently exceeded those of background cells, designating them as disease-relevant cells (**Methods**). While cells were individually scored, we also harnessed the knowledge of annotated cell types to facilitate biological interpretation (**Methods**).

#### Type 2 diabetes

There were 14 cell types identified in the human pancreas^28^ (**Fig. 4a**; **Methods**), among which two hormone-high cell types, namely GCG^high^ alpha cell and INS^high^ beta cell, were significantly enriched with scPRS-selected cells (adjusted *P* < 0.1, BH correction; **Fig. 4b,c**). The original study^35^ generating the pancreas snATAC-seq dataset had also linked INS^low^ beta cell, alongside INS^high^ beta cell, to T2D risk using the stratified linkage disequilibrium (LD) score regression (sLDSC)^36^. As another benchmark, we applied SCAVENGE^37^, a computational method that also enables single-cell-resolved cell prioritization, to the same data (**Methods**). In addition to GCG^high^ alpha cell and INS^high^ beta cell, SCAVENGE also identified GCG^low^ alpha cell (adjusted *P* < 0.1, BH correction; **Extended Data Fig. 3a**).

**Fig. 4.**
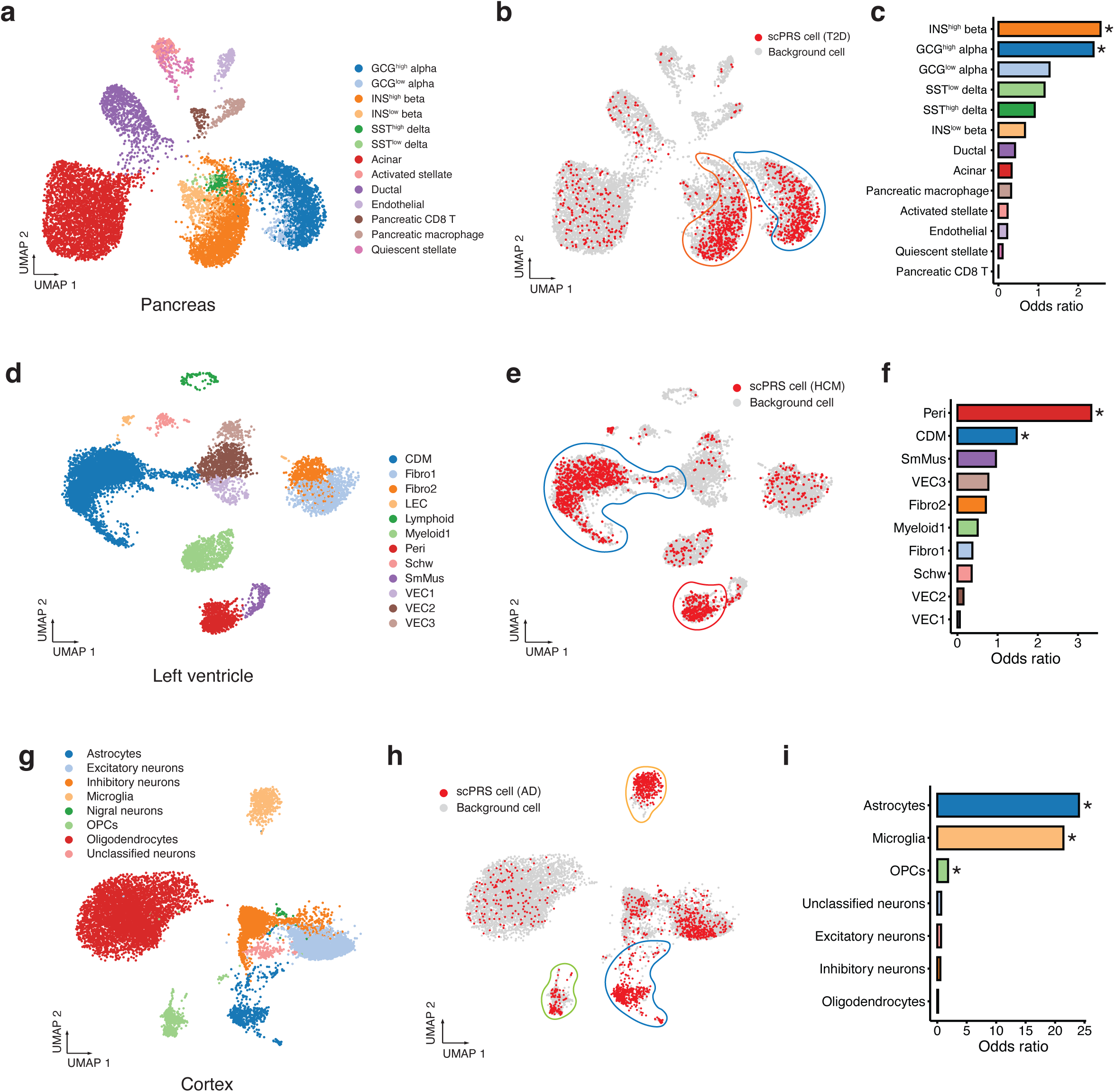
Disease-relevant cells identified by scPRS. **a**, The uniform manifold approximation and projection (UMAP) plot of the human pancreas snATAC-seq dataset. **b**, T2D-relevant cells prioritized by scPRS (in red). Cell clusters enriched with scPRS-prioritized cells are highlighted in closed curves with corresponding cell type colors. **c**, Enrichment of scPRS-selected T2D cells per cell type. *, adjusted *P* < 0.1. Odds ratio and *P*-value by one-sided Fisher’s exact test. **d**, UMAP plot of the human left ventricle snRNA-seq dataset. CDM, cardiomyocyte; Fibro, fibroblast; LEC, lymphatic endothelial cell; Peri, pericyte; Schw, Schwann cell; SmMus, smooth muscle cell; VEC, vascular endothelial cell. **e**, HCM-relevant cells prioritized by scPRS (in red). Cell clusters enriched with scPRS-prioritized cells are highlighted in closed curves with corresponding cell type colors. **f**, Enrichment of scPRS-selected HCM cells per cell type. *, adjusted *P* < 0.1. Odds ratio and *P*-value by one-sided Fisher’s exact test. **g**, UMAP plot of the human cortex scATAC-seq dataset. OPC, oligodendrocyte progenitor cell. **h**, AD-relevant cells prioritized by scPRS (in red). Cell clusters enriched with scPRS-prioritized cells are highlighted in closed curves with corresponding cell type colors. **i**, Enrichment of scPRS-selected AD cells per cell type. *, adjusted *P* < 0.1. Odds ratios and *P*-value by one-sided Fisher’s exact test. For robustness, small cell clusters with fewer than 150 cells were excluded from analysis and visualization in all diseases.

While pancreatic beta cell dysfunction and cell death are known as key processes in the development of T2D^38^, it is increasingly evident that T2D may result from defects in multiple cell types^39^. Notably, the alpha cell, regarded as the counterpart of the beta cell and responsible for producing glucagon, has been increasingly recognized as contributing to T2D pathogenesis^40,41,42^. Single-cell profiling further reveals the diversity within islet endocrine cells, spanning from fine-grained cell states to a continuous spectrum^35^. Our findings, coupled with prior research^35,43^, underscore the complexity of T2D pathogenesis involving multiple cell types within the pancreatic islets.

#### Hypertrophic cardiomyopathy

In the human left ventricle, a total of 17 cell types were identified (**Fig. 4d**; **Methods**). Among these, two cell types, including cardiomyocyte (CDM) and pericyte, presented significant enrichment with scPRS-selected cells (adjusted *P* < 0.1, BH correction; **Fig. 4e,f**). As benchmarking, we found no genetic enrichment within snATAC-seq peaks of all left ventricle cell types based on sLDSC (**Extended Data Fig. 3b**; **Methods**). SCAVENGE also linked cardiomyocytes to HCM (adjusted *P* < 0.1, BH correction), but no cell enrichment within pericytes was observed (**Extended Data Fig. 3c**). Cardiomyocytes, the primary cell type involved in the process of hypertrophy and thickening of heart muscle, play a pivotal role in HCM pathogenesis^44^. Pathogenic mutations disrupt the normal function of cardiomyocytes, leading to structural and functional abnormalities^44^, such as myocardial hypertrophy and fibrosis, contractile dysfunction, and arrhythmias. Our scPRS prediction not only reinforces the association between cardiomyocyte dysfunction and HCM but also extends this connection from protein function to noncoding gene regulation.

Cardiac pericytes interact with endothelial cells through both physical and paracrine mechanisms and are integral in maintaining cardiac and vascular homeostasis^45^. Despite being relatively understudied, the loss and dysfunction of pericytes have been associated with cardiomyopathy^46–48^. Our results agree with this connection and shed light on the potential causal involvement of pericytes in cardiac hypertrophy. Importantly, this link would not have been identified with either sLDSC or SCAVENGE.

#### Alzheimer’s disease

Eight major cell types were identified in the human cortex^30^ (middle frontal, superior and middle temporal gyri; **Fig. 4g**; **Methods**), among which three cell types were significantly enriched with scPRS-prioritized cells (adjusted *P* < 0.1, BH correction; **Fig. 4h,i**), including microglia, astrocyte, and oligodendrocyte progenitor cell (OPC). It is noteworthy that the original study^30^ generating the brain scATAC-seq dataset had linked only microglia to AD using sLDSC. Applying SCAVENGE to the same data revealed the same set of AD-relevant cell types as scPRS (adjusted *P* < 0.1, BH correction; **Extended Data Fig. 3d**).

The relationships between microglia and AD have been well established in previous literature^49^. Microglia play diverse roles, including immune response, phagocytosis, and synapsis modulation, contributing significantly to the development and progression of AD pathology. Moreover, genetic studies consistently prioritize microglia as the most prominent brain cell type associated with AD^50,51^. In recent years, accumulating evidence has underscored the essential role of astrocyte in AD pathogenesis through its reactivation or dysfunction^52,53^. Additionally, latest research has linked OPC to AD, likely due to its function in immune modulation and remyelination^54^. Our results reinforce these findings and offer further insights into the cellular heterogeneity of AD pathogenesis.

### scPRS reveals disease regulatory programs

As per the model design, scPRS prioritizes cells that contain disease-associated variants within their differentially accessible chromatin regions (**Fig. 1**). This feature empowers us to delve deeper into the regulatory circuits contributing upstream to the development of complex diseases across different cell types. To achieve this, we devised a layered multiomic strategy for systematically mapping cell-type-specific gene regulation underlying diseases (**Fig. 5a**; **Methods**).

**Fig. 5.**
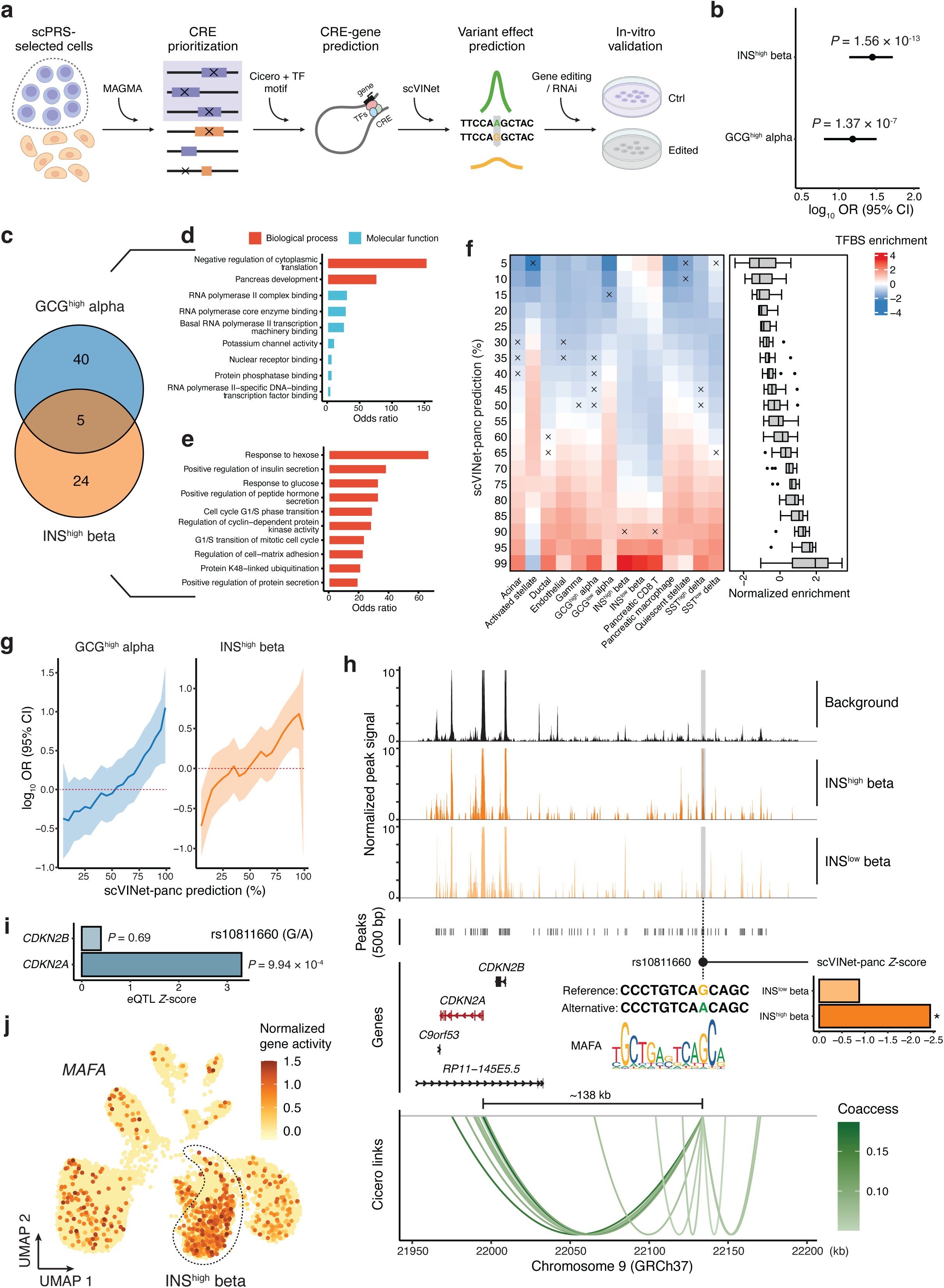
Cell-type-specific genetic regulation in T2D. **a**, Schematic of scPRS-based multiomic strategy for uncovering disease-relevant genetic regulation. CRE, cis-regulatory element; TF, transcription factor; RNAi, RNA interference; Ctrl, control. **b**, Enrichment of T2D-associated variants within CREs differentially accessible in scPRS-prioritized cells. Linkage disequilibrium (LD) threshold *r*^2^ = 0.1 was used in clumping to retrieve an independent variant set. *P*-value by two-sided Fisher’s exact test. OR, odds ratio; CI, confidence interval. The log_10_(OR) and 95% CI are annotated by the dot and error bar, respectively. **c-e**, Candidate T2D genes (**c**), and gene ontology (GO) enrichment for GCG^high^ alpha-cell genes (**d**) and INS^high^ beta-cell genes (**e**), respectively. Significant GO terms (adjusted *P* < 0.1, BH correction) with odds ratio greater than five are visualized. **f**, Enrichment of TF binding site (TFBS) disrupting variants within scVINet-panc-prioritized variants (various thresholds applied). Enrichment was estimated by *t*-statistics. The box plot center line, limits, and whiskers represent the median, quartiles, and 1.5x interquartile range (IQR), respectively. The dots indicate outliers falling above or below the end of the whiskers. ×, adjusted *P* > 0.1. **g**, Enrichment of scVINet-panc-prioritized T2D-associated variants (various thresholds applied) within T2D-CREs. OR and CI by two-sided Fisher’s exact test. OR is annotated by the solid line and 95% CI is represented by the shaded area. The red dashed line indicates null enrichment. **h**, Illustration of the genetic regulation of rs10811660 in INS^high^ beta cells. Bar plot: “*” indicates the percentage of scVINet-panc score is greater than 85%. Gene plot: the mapped target gene is highlighted in red. Link plot: links with coaccessibility greater than 0.05 are visualized; Coaccess, coaccessibility. **i**, Islet eQTL results for rs10811660. The effect allele is G. **j**, The uniform manifold approximation and projection (UMAP) plot of the pancreas snATAC-seq dataset showing the expression of *MAFA* in individual cells. Gene expression was estimated based on the gene activity computed by Signac^66^. INS^high^ beta cells are highlighted in the dashed closed curve.

For each disease-relevant cell type, we first identified candidate cis-regulatory elements (CREs) that were differentially accessible within scPRS-selected cells (**Methods**). Within these, we further prioritized CREs that were significantly enriched with disease-associated variants (referred to disease-relevant CREs) using MAGMA^55^ which is a region-based genetic association analysis approach (**Methods**). To chart the interactions between CREs and target genes, we performed coaccessibility analysis^56^ based on the single-cell ATAC-seq data (**Methods**), supplemented by the closest-gene strategy given its effectiveness in nominating disease genes^57^. For each cell type, this procedure yielded a set of candidate disease genes whose expression was associated by the prioritized CREs.

To fine-map candidate causal variants within disease-relevant CREs, we further developed a sequence-based deep learning model^58–60^, scVINet (**<u>s</u>**ingle-**<u>c</u>**ell-based **<u>v</u>**ariant **i**mpact **<u>net</u>**work), which predicts chromatin accessibility spanning various cell types solely based on the DNA sequence (**Extended Data Fig. 4a**; **Methods**). After training the model using the single-cell ATAC-seq data of a specific tissue (named scVINet-panc, scVINet-heat, and scVINet-brain for the pancreas, left ventricle, and cortex, respectively), we utilized in-silico mutagenesis to predict the functional effects of individual variants on chromatin accessibility across distinct cell types (**Extended Data Fig. 4a**; **Methods**). This completed the map of disease-relevant regulatory circuits composed of variant-CRE-gene trios. Follow-up experiments were carried out in corresponding cell types to validate our predictions.

#### Type 2 diabetes

We first observed a significant enrichment of T2D-associated variants (GWAS *P* < 5 × 10^−8^) within differentially accessible CREs of scPRS-prioritized cells (*P* < 1 × 10^−6^, two-sided Fisher’s exact test; **Fig. 5b** and **Extended Data Fig. 4b**; **Methods**). Using MAGMA, we identified 19 and 22 T2D-relevant CREs (referred to as T2D-CREs) in GCG^high^ alpha and INS^high^ beta cells, respectively (**Extended Data Fig. 4c** and **Supplementary Table 1**). Motif enrichment analysis for T2D-CREs uncovered transcription factors (TFs) of functional significance in corresponding cell types (**Extended Data Fig. 4c**; **Methods**). For example, TEAD1 is a critical beta-cell transcription factor necessary for coordinating various aspects of adult beta-cell function, including proliferative quiescence, mature identity, and functional competence to uphold glucose homeostasis^61,62^. MAFB, whose motif is enriched in both cell types, is another pivotal transcription factor in the islet. It is essential for the production and secretion of glucagon in alpha cells^63^, and also for the maturation of beta cells^64^. A recent study demonstrated that XBP1 plays a vital role in maintaining beta cell identity, repressing beta-to-alpha cell transdifferentiation, and is required for beta cell compensation and the prevention of diabetes in insulin resistance states^65^.

By mapping target genes of T2D-CREs, we identified 45 and 29 candidate risk genes in GCG^high^ alpha and INS^high^ beta cells, respectively (**Fig. 5c** and **Supplementary Table 1**). The function of alpha-cell genes was enriched with “Pancreas Development (GO:0031016)” and “RNA Polymerase Core Enzyme Binding (GO:0043175)” (adjusted *P* < 0.1, BH correction; **Fig. 5d**), while the function of beta-cell genes was enriched with “Response To Hexose (GO:0009746)”, “Positive Regulation Of Insulin Secretion (GO:0032024)”, and “Response To Glucose (GO:0009749)” (adjusted *P* < 0.1, BH correction; **Fig. 5e**). These results demonstrate the functional significance of our T2D risk genes in endocrine cells.

Trained on the pancreas snATAC-seq data, scVINet-panc exhibited high accuracy in peak prediction (AUROC: mean = 0.819, standard error (SE) = 0.011; AUPRC: mean = 0.639, SE = 0.044; **Extended Data Fig. 4d**). We validated our variant effect prediction using two different approaches: expression quantitative trait loci (eQTL) analysis and TF binding site prediction (**Methods**). Leveraging eQTL datasets generated in relevant tissues^67–70^, we observed that eQTLs tended to display larger effects based on scVINet-panc prediction in matching cell types compared to non-eQTLs (**Extended Data Fig. 4e**). Moreover, variants with larger effects were more likely to alter TF binding^71^ (**Fig. 5f**). These results indicate that scVINet-panc, while solely trained on the DNA sequence, had captured underlying gene regulatory mechanisms.

We also examined the functional effects of T2D-associated variants (GWAS *P* < 0.05) located within T2D-CREs in GCG^high^ alpha and INS^high^ beta cells (**Methods**). Interestingly, variants with larger predicted effect sizes were more enriched in T2D-CREs in corresponding cell types (**Fig. 5g**), providing an additional support for the functional importance of T2D-CREs we identified.

Combining multiomic evidence from eQTL, TF binding, and scVINet-panc prioritized T2D risk variants with functional implications (**Extended Data Fig. 4f**,g and **Supplementary Table 1**). One variant of particular interest is rs10811660, a T2D GWAS variant (GWAS *P* = 1.30 × 10^−11^, *β* = -0.13, effect/alternative allele is A)^27^ residing within an INS^high^-beta-cell-specific T2D-CRE (chr9:22,133,835-22,134,336; *P* = 1.91 × 10^−14^, log_2_ fold change (FC) = 4.99; **Fig. 5h**). Our prediction indicated that the alternative allele specifically decreased the CRE accessibility in INS^high^ beta cells (scVINet-panc INS^high^ beta cell *Z* = -2.43, percentile = 96.84%; **Fig. 5h**). Further, the affected CRE was found to be coaccessible with *CDKN2A* (coaccessibility = 0.159; **Fig. 5h**). Previous studies have demonstrated that the p16 inhibitor of cyclin-dependent kinase (p16^INK4A^), encoded by *CDKN2A*, restricts beta cell proliferation during aging, restricts beta cell regeneration, mediates overnutrition-related senescence, and reduces insulin secretory function^72^. While rs10811660 has also been linked to a *CDKN2A* paralog, *CDKN2B*, due to their distance proximity^72^, our coaccessibility analysis suggested that this association might be a false positive nomination (**Fig. 5h**). This conclusion was further supported by the islet eQTL data^70^, wherein rs10811660 was significantly associated with the expression of *CDKN2A* (*P* = 9.94 × 10^−4^, *Z* = 3.29) rather than that of *CDKN2B* (*P* > 0.05, *Z* = 0.40; **Fig. 5i**). Additionally, we found that the alternative allele A disrupted the binding motif of MAFA (*P* < 1 × 10^−4^, motifbreakR^73^; **Fig. 5h**; **Methods**), a critical regulator of pancreatic beta cell function^74^ which was specifically expressed in beta cells (**Fig. 5j**). Collectively, our analysis suggests a genetic regulation influencing T2D risk: the T2D risk allele G (rs10811660) increases the abundance of MAFA binding, which further up-regulates *CDKN2A* expression in INS^high^ beta cells. This aligns with previous evidence implicating that higher expression of *CDKN2A* may increase T2D risk^72^.

#### Hypertrophic cardiomyopathy

Following MAGMA and CRE-gene mapping, we identified 137 and 358 HCM-relevant CREs (referred to as HCM-CREs) linked to 199 and 492 target genes in cardiomyocytes (CDMs) and pericytes, respectively (**Supplementary Table 2**; **Methods**). Interestingly, we observed only minimal overlap, with just one CRE and 24 genes shared between these two cell types, highlighting their cell-type specificity.

Our motif enrichment analysis for HCM-CREs revealed transcription factors holding critical roles in corresponding cell types (**Fig. 6a**). For instance, TEAD1 is a pivotal regulator involved in maintaining the proper functioning of adult cardiomyocytes, whose loss of function has been associated with dilated cardiomyopathy (DCM)^75^. GATA4 exerts significant control over cardiac gene expression, impacting embryonic development, cardiomyocyte differentiation, and stress responsiveness of the adult heart^76^. NKX2-5 is a central regulator of heart development, and pathogenic mutations within it contribute to progressive cardiomyopathy and conduction defects^77^. Furthermore, RBPJ inactivation has been linked to the development of disease-promoting properties in brain pericytes^78^. STAT3 serves as a key regulator of cell-cell communication within the heart, a critical aspect of pericyte functionality^79^.

**Fig. 6.**
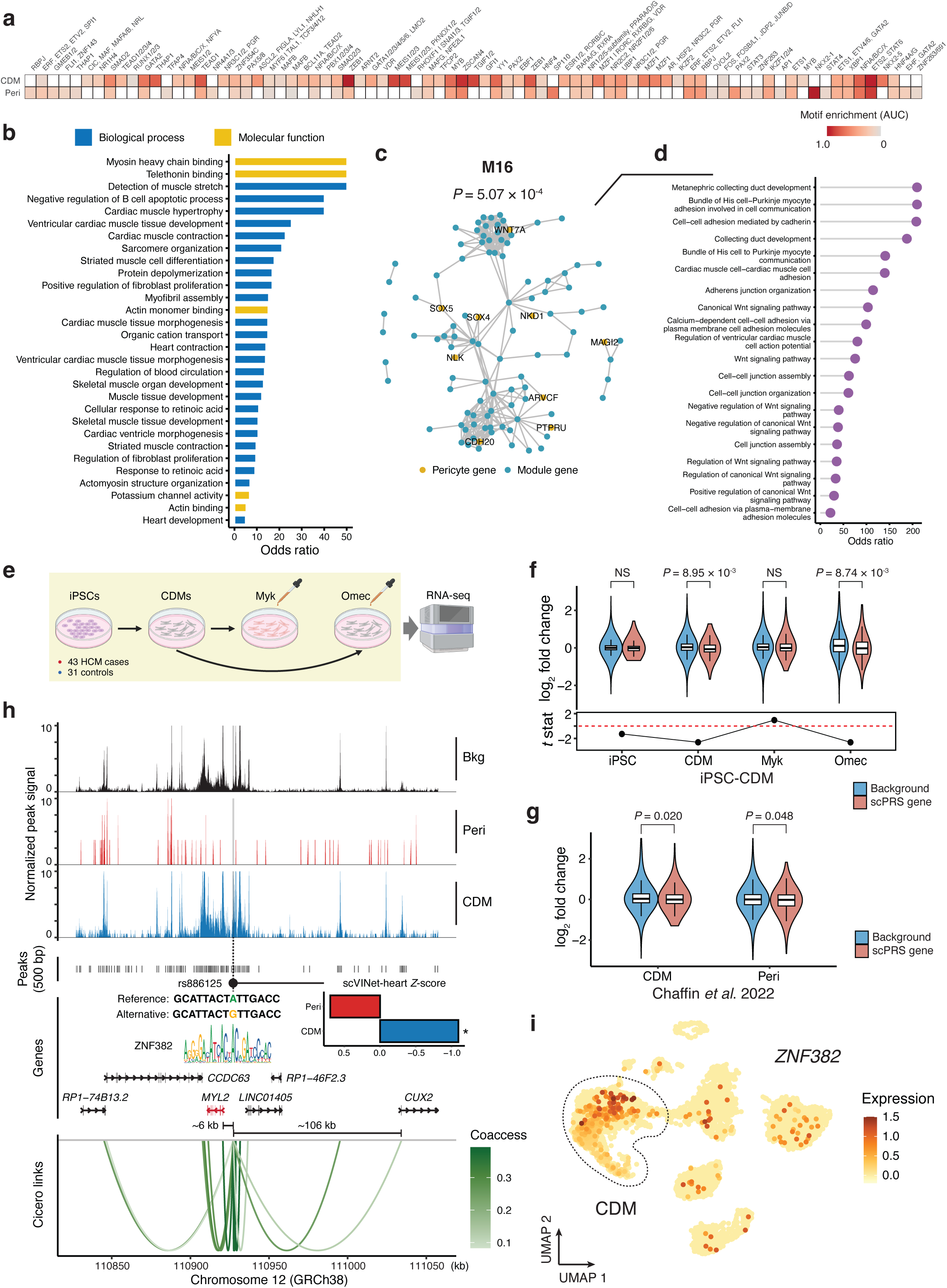
Cell-type-specific genetic regulation in HCM. **a**, Motif enrichment within HCM-CREs identified in two HCM-relevant cell types including cardiomyocyte and pericyte. Motif enrichment was measured by AUC. Row-wise standardization was performed. Only significant enrichment (adjusted *P* < 0.1, Bonferroni correction) is colored. CDM, cardiomyocyte; Peri, pericyte; AUC, the area under the receiver operating characteristic (ROC) curve. **b**, Bar plot of gene ontology (GO) enrichment for cardiomyocyte HCM risk genes. Significant GO terms (adjusted *P* < 0.1, BH correction) with odds ratio greater than five are shown. **c**, The network module M16 enriched with pericyte HCM risk genes. *P*-value by hypergeometric test. Edges between module genes are shown. **d**, Lollipop chart of GO enrichment (biological process) for M16 genes. Significant GO terms (adjusted *P* < 0.1, BH correction) are shown. **e**, Schematic of iPSC RNA-seq experiments. iPSC, induced pluripotent stem cell; Myk, Mavacamten; Omec, Omecamtiv mecarbil. **f**, Expression fold change comparison between HCM risk genes and the background transcriptome in cardiomyocytes across different conditions. The box plot center line, limits, and whiskers represent the median, quartiles, and 1.5x interquartile range (IQR), respectively. *P*-value by two-sided *t*-test. NS, not significant; stat, statistics. **g**, Expression fold change comparison between HCM risk genes and the background transcriptome in HCM-relevant cell types based on an HCM snRNA-seq study. *P*-value by two-sided *t*-test. **h**, Illustration of the genetic regulation of rs886125 in cardiomyocytes. Bar plot: “*” indicates scVINet-heart score percentage greater than 85%. Gene plot: differentially expressed target gene was mapped (in red). Bkg, background; Coaccess, coaccessibility. **i**, The uniform manifold approximation and projection (UMAP) plot of the left ventricle snRNA-seq dataset showing the expression of *ZNF382* in individual cells. Expression was estimated by normalized gene count. Cardiomyocytes are highlighted in the dashed closed curve.

HCM risk genes identified in cardiomyocytes exhibited functional significance in cardiomyocyte and cardiomyopathy, such as “Myosin Heavy Chain Binding (GO:0032036)”, “Cardiac Muscle Contraction (GO:0060048)”, and “Sarcomere Organization (GO:0045214)” (adjusted *P* < 0.1, BH correction; **Fig. 6b**). No GO enrichment was observed for pericyte genes, suggesting a significant functional diversity within this gene set. To better dissect this heterogeneity, we carried out a network analysis based on protein-protein interactions (PPIs)^80^ (**Methods**). Notably, one network module M16 was significantly enriched with our HCM pericyte genes (*P* = 5.07 × 10^−4^, hypergeometric test; adjusted *P* = 0.034, BH correction; **Fig. 6c**). This module of genes displayed GO enrichment in various pericyte functions, such as “Cell-Cell Adhesion Mediated By Cadherin (GO:0044331)”, “Cell-Cell Junction Assembly (GO:0007043)”, and “Cadherin Binding (GO:0045296)” (adjusted *P* < 0.1, BH correction; **Fig. 6d** and **Extended Data Fig. 5a**).

To gain a deeper understanding of the function of our HCM genes in the disease context, we analyzed an RNA-seq dataset^24^ of induced pluripotent stem cell (iPSC)-derived cardiomyocytes (iCDMs) obtained from 43 HCM cases and 31 healthy controls (**Fig. 6e**). Bulk RNA-seq profiling was conducted under four conditions: iPSC, differentiated cardiomyocyte, Mavacamten^81^ (an HCM drug recently approved by FDA) treated cardiomyocyte, and Omecamtiv mecarbil^82^ (a heart failure drug serving as the negative control) treated cardiomyocyte. Notably, although our cardiomyocyte HCM genes exhibited no expression difference in iPSCs between HCM cases and healthy controls, their expression was significantly reduced in differentiated HCM cardiomyocytes compared to control cells (*P* = 8..95 × 10^−3^, two-sided *t*-test; **Fig. 6f** and **Supplementary Table 3**; **Methods**). Intriguingly, the down-regulation of our HCM genes was rescued by Mavacamten treatment (*P* = 0.017, two-sided *t*-test) but persisted in Omecamtiv mecarbil treatment (*P* > 0.05, two-sided *t*-test; **Fig. 6f**). The reduced expression of HCM genes identified in cardiomyocytes and pericytes was also confirmed in corresponding cell types based on an independent HCM single-cell transcriptome dataset^83^ (cardiomyocyte: *P* = 0.02; pericyte: *P* = 0.048, two-sided *t*-test; **Fig. 6g** and **Extended Data Fig. 5b**,c). Taken together, these results provide strong evidence demonstrating the disease relevance of our HCM genes.

We trained scVINet-heart based on the snATAC-seq dataset of the left ventricle (AUROC: mean = 0.846, SE = 0.019; AUPRC: mean = 0.658, SE = 0.032; **Extended Data Fig. 5d**). The variant effects predicted by scVINet-heart agreed well with eQTL profiling^67,84^ and TF binding site prediction (**Extended Data Fig. 5e**,f). Moreover, HCM-CREs within corresponding cell types presented increased enrichment with larger-effect HCM-associated variants (GWAS *P* < 0.05; **Extended Data Fig. 5g**), highlighting their functional significance.

Our scVINet-heart-based variant effect prediction, together with eQTL and TFBS analyses, prioritized novel cell-type-specific HCM risk variants (**Extended Data Fig. 5h**,i and **Supplementary Table 2**). As an example, the cardiomyocyte-specific HCM-CRE (chr12:110,927,025-110,927,526; *P* = 2.5 × 10^−3^, log_2_ FC = 1.94; **Fig. 6h**) contains a nominally significant GWAS variant rs886125 (GWAS *P* = 0.019, *β* = -0.149, effect/alternative allele = G)^26^, and was coaccessible (coaccessibility = 0.367) with *MYL2* which is a widely recognized HCM risk gene^61^. Based on our scVINet-heart prediction, the alternative allele of rs886125 specifically diminished CRE accessibility within cardiomyocytes (scVINet CDM *Z* = -1.10, percentile = 87.62%; **Fig. 6h**). We further predicted that it disrupted the binding site of a transcription factor ZNF382 (*P* < 1 × 10^−4^, motifbreakR; **Fig. 6h**), which is known as a transcriptional repressor^85^. These results suggest that the risk-increasing allele A, bound by ZNF382, would lower the expression of *MYL2* in cardiomyocytes. This was reinforced by the eQTL data^68^ in which the alternative allele G was associated with increased expression of *MYL2* (*P* = 0.011, *β* = 0.125; GTEx artery aorta). Notably, using our paired single-nucleus RNA sequencing (snRNA-seq) data, we observed that *ZNF382* was specifically expressed in cardiomyocytes (**Fig. 6i**), highlighting its cell-type-specific role in gene regulation.

#### Alzheimer’s disease

We first confirmed a significant enrichment of AD-associated variants (GWAS *P* < 5 × 10^−8^) within differentially accessible CREs in scPRS-prioritized cells (*P* < 5 × 10^−3^, two-sided Fisher’s exact test; **Extended Data Fig. 6a**). We identified 39, 57, and 6 AD-relevant CREs (referred to as AD-CREs), which were linked to 71, 118, and 33 target genes in astrocytes, microglia, and OPCs, respectively (**Fig. 7a** and **Supplementary Table 4**; **Methods**). Interestingly, numerous AD-CREs and genes were shared across different cell types, among which we recognized multiple well-established AD genes, such as the *APOE* region genes (*BCAM*, *NECTIN2*, *TOMM40*, *APOE*, and *APOC1*), *BCL3*, and *PPP1R37*. This reinforced the link between these genes and AD and signified their versatile roles in disease pathogenesis.

**Fig. 7.**
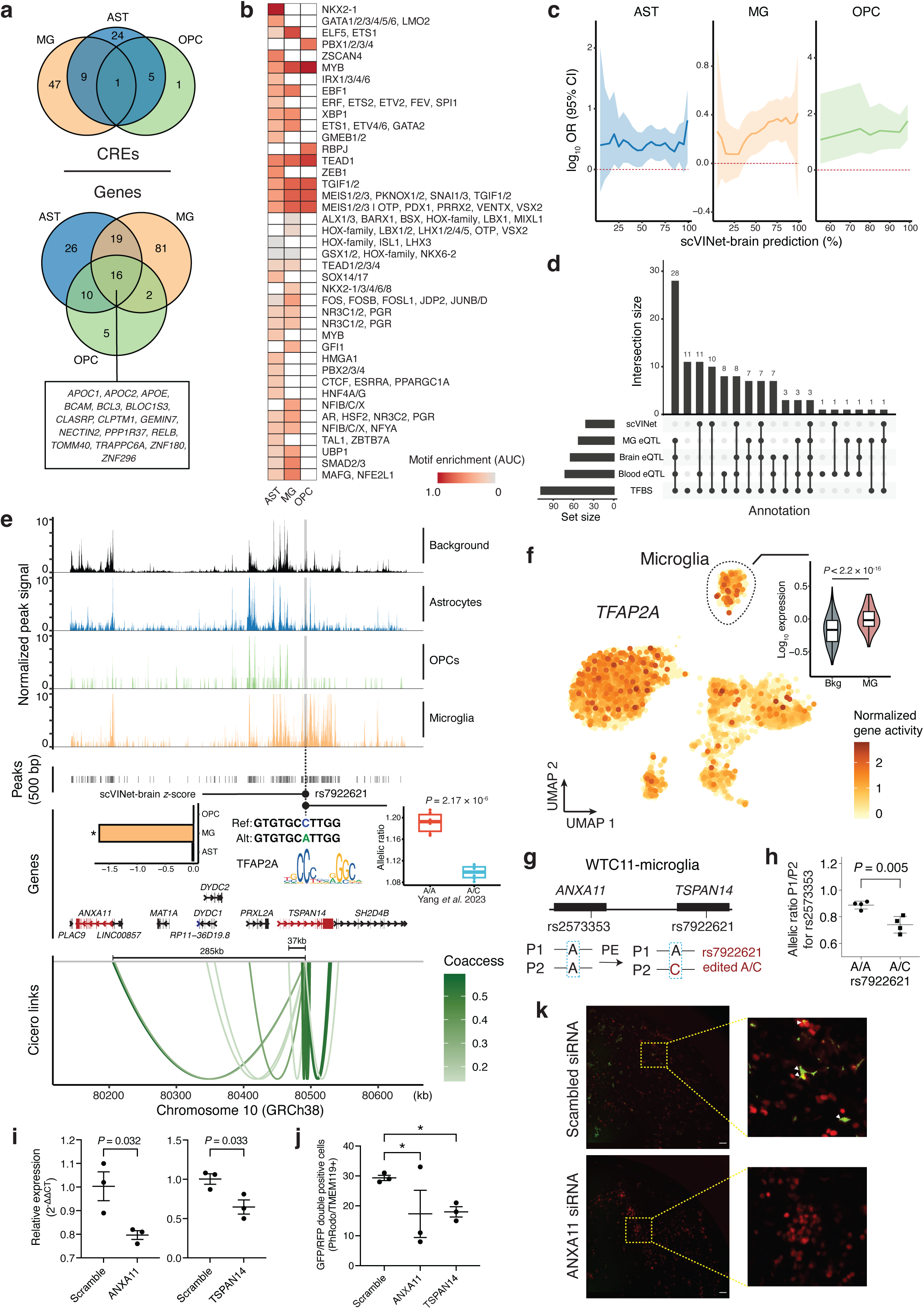
Cell-type-specific genetic regulation in AD. **a**, Venn diagram of AD-relevant CREs (top) and genes (bottom) identified by our multiomic strategy. AST, astrocyte; MG, microglia; OPC, oligodendrocyte progenitor cell. **b**, Motif enrichment within AD-CREs identified in different cell types. Motif enrichment was measured by AUC. Column-wise standardization was performed. Only significant enrichment (adjusted *P* < 0.1, Bonferroni correction) is colored. AUC, the area under the receiver operating characteristic (ROC) curve. **c**, Enrichment of scVINet-brain-prioritized AD-associated variants (various thresholds applied) within AD-CREs. OR and CI by two-sided Fisher’s exact test. OR is annotated by the solid line and 95% CI is represented by the shaded area. Red dashed line indicates null enrichment. OR, odds ratio; CI, confidence interval. **d**, Summary statistics of fine-mapped AD risk variants in microglia using different annotations. TFBS, transcription factor binding site. **e**, Illustration of the genetic regulation of rs7922621 in microglia. Box plot: the box plot center line, limits, and whiskers represent the median, quartiles, and 1.5x interquartile range (IQR), respectively; *P*-value by two-sided *t*-test. Variant plot: ref, reference; alt, alternative. Bar plot: “*” indicates scVINet-brain score percentage greater than 85%. Gene plot: differentially expressed target genes were mapped (in red). Link plot: links with coaccessibility greater than 0.05 are shown; Coaccess, coaccessibility. **f**, The uniform manifold approximation and projection (UMAP) plot of the cortex scATAC-seq dataset showing the expression of *TFAP2A* in individual cells. Violin plot: *P*-value by two-sided *t*-test; Bkg, background. Gene expression was estimated based on the gene activity computed by Signac. Microglia are highlighted in the dashed closed curve. **g**, Diagram showing the haplotypes of variants in wide type (WT) and rs7922621 prime edited WTC11-derived microglia. The P1 allele has the risk allele (A), while the P2 allele has the non-risk allele (C). PE, prime editing. **h**, Allelic imbalance between P1 and P2 alleles for *ANXA11* quantified by rs2573353 in rs7922621 wild-type (A/A) and prime-edited (A/C) WTC-derived microglia (*n* = 4 replicates). The center line and error bar represent the mean and standard deviation, respectively. *P*-value two-sided *t*-test. **i**, RT-qPCR quantification of relative mRNA levels in iMGs treated with siRNAs targeting AD genes or scrambled siRNA (*n* = 3 replicates). mRNA levels were normalized to GAPDH. *P*-value by two-sided *t*-test. The center line and error bar indicate the mean and standard error, respectively. RT-qPCR, reverse transcription-quantitative polymerase chain reaction; iMG, iPSC-derived microglia-like cell. **j**, Quantification of the number of TMEM119+ cells colocalized with pHrodo particles indicating phagocytosed beads (*n* = 3 replicates). One-way ANOVA was used for comparison between siRNA targeting AD genes and scrambled siRNA. The center line and error bar represent the mean and standard error, respectively. *, adjusted *P* < 0.05 by the Benjamini-Hochberg (BH) correction. **k**. Representative images of TMEM119+ (red) iMGs treated with *ANXA11* siRNA or scrambled siRNA showing colocalization of phagocytosed pHrodo particles (green, highlighted with arrows). Images were captured two hours after incubation with pHrodo. Part of the images are zoomed in for better visualization. siRNA, small interfering RNA. Scale bar = 100μm.

Next, we examined the function of AD-CREs and candidate genes in corresponding cell types. In particular, we found that AD-CREs were enriched with binding motifs of cell-type-critical transcription factors (**Fig. 7b**). For example, astrocyte AD-CREs displayed exclusive motif enrichment for GATA4, a regulator of astrocyte cell proliferation and apoptosis^86^; microglia AD-CREs exhibited significant motif enrichment for SMAD3, which cooperates with PU.1 to enable transcription of some microglia-specific genes^87^; OPC AD-CREs were exclusively enriched with the motif of RBPJ, which is a repressor of a major determinant of oligodendrocyte differentiation and myelination – OLIG2^88^. Moreover, our AD candidate genes also presented cell-type-specific functions. For instance, astrocyte AD genes were enriched with the function of “Regulation Of Complement Activation, Classical Pathway (GO:0030450)”, microglia AD genes displayed enrichment in “Negative Regulation Of Endocytosis (GO:0045806)”, and OPC AD genes exhibited significant enrichment in “IκB kinase/NF-κB Signaling (GO:0007249)” (adjusted *P* < 0.1, BH correction; **Supplementary Table 5**).

To characterize the function of variants within AD-CREs, we trained scVINet-brain based on the cortex scATAC-seq data (AUROC: mean = 0.916, SE = 0.017; AUPRC: mean = 0.795, SE = 0.059; **Extended Data Fig. 6b**). We confirmed the agreement in variant effect inference between scVINet-brain and two other approaches, including QTL (expression and chromatin accessibility) analysis and TF binding site prediction (**Extended Data Fig. 6c**,d). Based on scVINet-brain, we uncovered an enrichment of large-effect AD-associated variants (GWAS *P* < 0.05) within AD-CREs in all three AD-relevant cell types, wherein the enrichment was positively correlated with variant effect size (**Fig. 7c**).

We successfully fine-mapped AD risk variants by combining scVINet-brain, QTL, and TF binding site prediction (**Fig. 7d** and **Extended Data Fig. 6e**,f). Among the prioritized variants, we recognized numerous cell-type-specific risk loci that were previously reported in the literature. For example, the AD risk variant rs10792832 (GWAS *P* = 7.56 × 10^−16^, *β* = -0.12, effect allele/reference = A)^25^ has been associated with the deactivation of a microglia-specific CRE for *PICALM*^51^, aligning with our prediction (scVINet-brain microglia *Z* = -1.98, *PICALM* coaccessibility = 0.246; **Supplementary Table 4**). Another AD risk variant rs13025717 (GWAS *P* = 2.98 × 10^−15^, *β* = 0.13, effect/alternative allele = T), which represses a microglia CRE of *BIN1*^30^, was also prioritized by our prediction (scVINet-brain microglia *Z* = -2.60, *BIN1* coaccessibility = 0.382; **Supplementary Table 4**). Moreover, a recent study validated the role of rs1532278 (GWAS *P* = 3.27 × 10^−16^, *β* = -0.13, effect/reference allele = T) in modulating *CLU* expression in astrocytes^89^, providing support for our prediction (scVINet-brain astrocyte *Z* = -0.498, *CLU* coaccessibility = 0.356; **Supplementary Table 4**).

In addition to known AD risk loci and genes, our analysis also discovered novel genetic factors. One of particular interest is rs7922621 which is nominally significant genome-wide (GWAS *P* = 2.78 × 10^−5^, *β* = 0.08, effect/alternative allele = A)^25^. This variant resides within a microglia-specific AD-CRE (chr10:82,251,479-82,251,979; *P* = 1.99 × 10^−19^, log_2_ FC = 2.39; **Fig. 7e**). According to the scVINet-brain prediction, rs7922621 diminished the accessibility of its surrounding CRE exclusively in microglia but not in other cell types (scVINet-brain microglia *Z* = -1.68, percentile = 96.63%; **Fig. 7e**). Coaccessibility analysis further predicted that this CRE regulates the expression of two genes: *ANXA11* and *TSPAN14* (**Fig. 7e**). Importantly, a recent study reported a reduction in local chromatin accessibility associated with rs7922621 in human pluripotent stem cell (hPSC)-derived microglia^90^. They further validated the reduced expression of *TSPAN14* due to rs7922621 using prime editing (*P* = 2.17 × 10^−6^, two-sided *t*-test; **Fig. 7e**). Of note, another variant, rs7910643, located within the same CRE and in strong LD with rs7922621 (*r*^2^ = 1.0, estimated in the 1000 Genomes European population), was shown to be non-functional^90^, consistent with our prediction (scVINet-brain microglia *Z* = 0.29, percentile < 85%; **Supplementary Table 4**).

To further elucidate the regulatory program involving rs7922621, we conducted TF motif analysis and identified one transcription factor, TFAP2A, whose binding site was disrupted by rs7922621 (*P* < 1 × 10^−4^, motifbreakR; **Fig. 7e**). TFAP2A belongs to the transcription factor family activator protein 2 (TFAP2) family, known for its pivotal role in regulating both embryonic and oncogenic development^91^. Importantly, *TFAP2A* expression showed a significant elevation in microglia compared to other cell types (*P* < 2.2 × 10^−16^, two-sided *t*-test; **Fig. 7f**), suggesting its functional significance in microglia, though further evidence is required to validate these conclusions.

### Prime editing of rs7922621 alters expression of both ANXA11 and TSPAN14 in microglia

Our scPRS-based analysis pinpointed rs7922621 (chr10:82,251,544:C>A) as a candidate AD risk variant and predicted that it regulates two genes including *ANXA11* and *TSPAN14* through altering the accessibility of a microglia-specific CRE (chr10:82,251,479-82,251,979; **Fig. 7e**). In a prior study^90^, Yang et al. validated the association between rs7922621 and reduced CRE, and further demonstrated that the prime editing (PE) of rs7922621, converting the risk allele (A) to the non-risk allele (C) in WTC11 (A/A to A/C)-derived microglia (a male iPSC line), led to an increase in *TSPAN14* expression. Leveraging the rs7922621-edited clones^90^, we further examined its regulatory role on *ANXA11* (**Fig. 7g**; **Methods**). Surprisingly, we observed a similar trend in the allelic expression changes of *ANXA11* associated with rs7922621 in WTC11-derived microglia, with the edited non-risk allele upregulating *ANXA11* compared to the risk allele (*P* = 0.005, two-sided *t*-test; **Fig. 7h**). We note that in contrast to *TSPAN14*, *ANXA11* is in a long-range interaction (∼285 kb) with rs7922621 (**Fig. 7e**). Altogether, these results suggest that rs7922621 regulates both *ANXA11* and *TSPAN14* in microglia, with the AD risk allele (A) reducing their expression.

### Suppression of ANXA11 and TSPAN14 impairs microglia phagocytosis

To further elucidate the function of *ANXA11* and *TSPAN14* in microglia, we examined the effect of knockdown of these two genes on microglial phagocytic activity. In particular, we individually suppressed *ANXA11* and *TSPAN14* in iPSC-derived microglia-like cells (iMGs)^92,93^ using small interfering RNA (siRNA), and then measured phagocytosis activity using a fluorescent read-out of pHrodo particles (**Methods**). Expression reduction using siRNA treatment was confirmed for both genes (*P* < 0.05, two-sided *t*-test; **Fig.7i**). Remarkably, suppression of these two genes both resulted in decreased iMG uptake of pHrodo particles compared to the scrambled siRNA treatment (adjusted *P* < 0.05, one-way ANOVA with BH correction; **Fig. 7j,k**). Our results demonstrate the functional significance of *ANXA11* and *TSPAN14* whose suppression impairs microglia phagocytosis. This further supports the pivotal role of rs7922621 in impacting AD risk by modulating microglia function.

## Discussion

Genome-wide association studies have significantly advanced our understanding of the genetic basis of complex human diseases^94^. Traditionally, these studies primarily aim to identify genetic loci that reach genome-wide significance (i.e., GWAS *P* < 5 × 10^−8^) to reduce the risk of false-positive findings. However, for many diseases, the best prediction performance is achieved by including nominally significant or even non-significant genetic variants when constructing PRS^95^. This suggests that the genetic factors contributing to diseases extend beyond those genome-wide significant loci and cannot be fully discovered by conventional approaches^96^. While scientists have been calling for larger GWAS consortia and meta-analysis to discover more disease risk loci^97^, it remains an open question how to increase the discovery power given relatively limited sample size. Incorporating prior knowledge or multiomic data into genetic association analysis has been proven as an effective solution^98,99^.

PRS has been demonstrated as a powerful tool for predicting an individual’s disease risk. However, it lacks the ability to provide insights into disease mechanisms. From the perspective of modern machine learning, model interpretation is critical in uncovering latent features that contribute to prediction and understanding how models make decisions^100^. As a score computed by aggregating a wide range of variants, PRS offers limited knowledge of the significance of each variant. Moreover, distinguishing causal variants from statistically correlated elements poses an even greater challenge. For example, a variant can be associated with the disease through its linkage with another causal variant, yet both are treated equivalently within a PRS model. This lack in biology-informed model interpretability can, in turn, constrain prediction performance such as generalizability^101^.

We have designed scPRS, a geometric deep learning-based PRS to address these challenges. scPRS leverages single-cell epigenetic data to dissect the genome-wide PRS and then integrates single-cell-level PRSs using a graph neural network. By breaking down PRS into high-resolution components informed by cell functions, scPRS not only enhances its prediction power but also allows for a systematic exploration of cellular and molecular disease basis. Applications to various diseases have shown that scPRS outperformed a variety of conventional PRS methods. Importantly, this superior prediction performance of scPRS was achieved by only utilizing less than 10% of all the variants (i.e., those variants located within open chromatin regions), highlighting the importance of incorporating functional data^101^ and suggesting a significant contribution of noncoding variants to disease risk^102^.

We have showcased the effectiveness of scPRS in identifying disease-critical cells. Our method is not confined to pre-clustering and annotating single-cell data, offering an unbiased analysis and the potential to discover novel disease-critical cell populations. Several recent studies^37,103,104^ have also achieved prioritization of disease-relevant cells at the single-cell level. However, these approaches are designed relying on GWAS summary statistics and thus lack prediction power. Moreover, superior to these methods, scPRS has been demonstrated to facilitate pinpointing disease risk variants, genes, and regulatory programs across different cellular contexts, substantially enhancing the power and resolution of genetic discovery. This advancement is exemplified by rs7922621 which was pinpointed by scPRS-based analysis as a candidate AD risk variant while missed by GWAS due to its nominal significance. rs7922621 has also been nominated by two latest studies^90,105^, wherein it was mapped to *TSPAN14* in microglia as the target gene. Our scPRS analysis further linked rs7922621 to another gene *ANXA11*. *ANXA11* is known as a risk factor for amyotrophic lateral sclerosis (ALS) and it mediates neuronal RNA transport by tethering RNA granules to actively-transported lysosomes^106^. Our results suggest a new role of *ANXA11* in microglia and AD pathology. We have experimentally validated the regulatory relationship between rs7922621 and *ANXA11*, as well as the function of these two genes (*ANXA11* and *TSPAN14*) in maintaining microglia phagocytosis. Our data support the model that rs7922621 increases AD risk by reducing a microglia CRE targeting *ANXA11* and *TSPAN14* and then suppressing their expression, which further impairs microglia phagocytosis.

The genetic study of hypertrophic cardiomyopathy has been traditionally focused on rare pathogenic coding variants^44^. However, approximately 40% of HCM patients remained unexplained by known pathogenic variants^44^. Previous HCM GWASs for common variants have been underpowered, likely due to the limited number of patients recruited, resulting in an incomplete knowledge of the genetic architecture^107^. Our scPRS-based analysis significantly expands our understanding of HCM genetics, highlighting the critical role of common noncoding variants in influencing HCM risk. Our findings underscore the importance of regulatory variants that have been largely overlooked in the HCM field. These variants act as risk modifiers through modulating the expression of their target genes, including known HCM risk genes such as *MYL2*. Although further validations are necessary, our results shed light on the complexity of HCM genetics and biology.

Single-cell genetics is an emerging field that is reshaping our understanding of genotype-phenotype relationships^15^. By integrating single-cell genomic data into genetic analysis, single-cell genetics provides a novel instrument to connect genetic variants with diverse cellular processes. This is well exemplified by single-cell eQTL (sc-eQTL) studies^108–110^ which enable the identification of context-dependent eQTLs that vary with cell states or cell types. Our scPRS lays the methodological foundation of single-cell genetics and represents a pioneering step toward unveiling the genetic basis of complex diseases within a single-cell-resolved context.

Following the same design principle, scPRS could be extended to incorporate additional omic data, such as single-cell RNA-seq data^111,112^, single-cell DNA methylation data^113,114^, and even rare variants^115,116^. In summary, scPRS stands as a next-generation framework for simultaneous disease prediction and biological discovery, enabling dissection of the genetic, cellular, and molecular heterogeneity underlying complex diseases.

## Data availability

The PBMC multiome dataset is available from 10x Genomics (https://support.10xgenomics.com/single-cell-multiome-atac-gex/datasets/1.0.0/pbmc_granulocy <u>te_sorted_10k</u>). The single-cell multiome (snRNA-seq and snATAC-seq) data of the human left ventricle are publicly accessible through ENCODE 4 (https://www.encodeproject.org/multiomics-series/ENCSR245BNT/). All other single-cell datasets were obtained from their original publications^28,30^. The genotype data used in simulation are available from ref. ^20^. The whole-genome sequencing and iPSC RNA-seq data for HCM are available from ref. ^24^. Individual-level genotype-phenotype data for T2D and AD were sourced from the UK Biobank (Application ID 41751). The HCM snRNA-seq dataset was obtained from ref. ^83^. All GWAS summary statistics datasets were acquired from their original publications^19,25–27^. The GTEx and islet eQTL datasets were downloaded from the eQTL Catalogue (https://www.ebi.ac.uk/eqtl/). Other eQTL and caQTL datasets were obtained from the original publications^51,67,70,117^.

## Code availability

The scPRS software is available at https://github.com/szhang1112/scPRS. We implemented scVINet based on Selene v0.5.0 (https://github.com/FunctionLab/selene). Baseline PRS implementations include C+T (PLINK v1.9), LDpred2 (bigsnpr v1.12.2), and Lassosum v0.4.5. Logistic regression models were implemented using Scikit-learn v1.2.2. We also implemented SCAVENGE (https://github.com/sankaranlab/SCAVENGE) and stratified LDSC v1.0.1 (https://github.com/bulik/ldsc) for disease cell prioritization. Single-cell data processing and analysis were performed based on Seurat v4.3.0, Scanpy v1.8.1, SnapATAC2 v2.6.0, Signac v1.9.0, and ALLCools v1.0.22 (https://github.com/lhqing/ALLCools). GO analysis was conducted using Enrichr (https://maayanlab.cloud/Enrichr/) if not specified. TF motif analysis was carried out using GimmeMotifs v0.18.0 (https://gimmemotifs.readthedocs.io/en/master/). We performed TF binding site prediction using SNP2TFBS (https://epd.expasy.org/snp2tfbs/) and motifbreakR v2.15.5 (https://github.com/Simon-Coetzee/motifBreakR). All statistical analyses were performed using Python v3, R v4, and Prism v10.

## Acknowledgements

We thank Dr. Ryan Corces for providing fragment files for the human brain scATAC-seq data and for his detailed responses to our questions regarding this dataset. This work was supported by NIH (CEGS 5P50HG007735 to M.P.S., R01AG079291 and RF1AG079557 to Y.S., 2R01NS097850 and 1R01NS131409 to J.K.I.). This work was also supported by the Tau Consortium and John Douglas French Alzheimer’s Foundation (to J.K.I.). J.K.I. is the John Douglas French Alzheimer’s Foundation Endowed Associate Professor of Stem Cell Biology and Regenerative Medicine. Figures 1,5a, 6e and Extended Data Figure 4a were created using BioRender.

## Author contributions

S.Z. conceived and designed the study. S.Z. and H.S. developed and implemented scPRS and scVINet. X.Y. and Y.L. (supervised by Y.S.) conducted prime editing experiments. J.R.-S. and S.T. (supervised by J.K.I.) performed siRNA and pHrodo phagocytosis experiments. S.Z., H.S., J.Zhou analyzed the data with assistance from J.R.-S., X.Y., Y.L., J.C.-K., E.M. and C.Z. All authors were responsible for data interpretation. S.Z., J.Zeng, P.S.T. and M.P.S. supervised the project. S.Z., H.S., J.Zhou, J.Zeng, P.S.T. and M.P.S. drafted the manuscript with assistance from all other authors. All authors meet the four ICMJE authorship criteria and were responsible for approving the final version for publication and for accuracy and integrity of the work.

## Methods

### Single-cell multiome dataset

Single-cell multiome (snRNA-seq + snATAC-seq) data of the human left ventricle were processed and clustered based on the RNA modality using Scanpy^118^. The cells with high-quality RNA information (total detected gene > 500, total unique molecular identifier (UMI) < 20000, and mitochondrial read percentage < 10%) were selected for further analysis. Doublets were filtered using scrublet^119^ with parameters min_counts=1, min_cells=10, min_gene_variability_pctl=90, and n_prin_comps=30. The thresholds for doublet removal were decided per sample based on the distribution of doublet scores in real versus simulated cells. Top 3000 highly variable genes were selected by combining the results from each sample separately with seurat_v3 mode. The cell-by-gene count matrices were normalized and scaled. ALLCools with a Python implementation of Seurat integration was used for correction of batch effect between samples with 50 principle components (PCs) and 30 canonical correlation dimensions^114,120^. Leiden clustering was performed on a *k*-nearest neighbor (kNN; *k* = 25) graph. The cell clusters were annotated and merged to cell types by comparing the expression level of predefined marker genes across clusters. The marker genes in Litviňuková et al. 2020 and Tucker et al. 2020^121,122^ were used to annotate the heart cell types.

We also examined the ATAC modality of these cells following the methods described below to ensure these cells also have high-quality open chromatin information. The cells that did not pass ATAC quality controls (QCs) or constituted an ambiguous cluster in ATAC cell embedding were removed, resulting in 10,233 cells retained for downstream analysis.

### Single-cell ATAC-seq datasets

The cell type labels for the human pancreas and cortex in the original datasets^28,30^ were used. To generate cell embeddings, single-cell ATAC-seq data were processed and clustered using snapATAC2^123^ and ALLCools^114,120^. The fragment files were processed to generate cell-by-bin matrices at 5 kb resolution using snapATAC2^124^. The cells with 2000 < total reads < 50000 and transcription start site (TSS) enrichment > 5 or 7 based on the distribution in specific samples were retained. The cell embeddings were computed with latent semantic analysis (LSI) and batch effects were corrected using the CCA-LSI mode in ALLCools. Cell-by-peak matrices at 500-bp resolution were generated by calling peaks per cell cluster using snapATAC2. For cortex data, superior and middle temporal gyri and middle frontal gyrus samples were used for AD analysis, resulting in 11,738 cells. For pancreas data, we randomly sampled 10,000 (out of 64,948) cells covering all annotated cell types for computational acceleration.

### Cell-cell similarity network

Following the previous study^125^, we utilized the mutual *k*-nearest neighbors (M-kNN) to measure the similarity between two different cells. We first employed latent semantic indexing (LSI) to extract low-dimensional embeddings for individual cells. For cortex and left ventricle datasets encompassing multiple samples, batch effects were corrected using both canonical correlation analysis (CCA) and Harmony^126^, and integrated latent embeddings were adopted. Next, we computed the Euclidean distance for pairs of cells using their embeddings, and then constructed the kNN graph 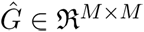 based on this distance matrix, in which we defined 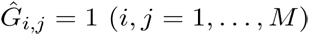 if cell *j* is within the top *k* closest cells of cell *i*, and 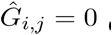 otherwise. The M-kNN graph *G* was then defined as a graph whose edges link nodes (i.e., cells) that are mutually *k* nearest neighbors of each other, and was calculated by 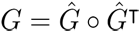 where O denotes the element-wise multiplication.

### Target genotype cohorts

T2D and AD target cohorts were constructed based on the UK Biobank (UKBB). All the disease cases were defined based on the ICD-10 (10th revision of the International Statistical Classification of Diseases and Related Health Problems) code. In particular, all Caucasian individuals with a disease ICD-10 code in inpatient record, death record, or diagnosis summary record were defined as the disease patients. We used E11.9 and G30.9 for AD and T2D, respectively. This resulted in 1,096 T2D and 932 AD cases. We randomly sampled an equal number of healthy controls by matching sex, age, and ancestry information for each case group. In addition, individuals with a similar or related phenotype with the disease (T2D: E10, E11, E12, E13, E14, E23.2, N08.3, N25.1, O24, P70.2, Z13.1, Z83.3, R73.9; AD: F00, G30, F01, F02, F03, F05) were excluded from constructing the control group. In this study, overweight individuals (body mass index (BMI) >= 25) were excluded from constructing the T2D cohort. BMI for each individual was defined as the mean of four BMI measurements in the UKBB Data-Field 21001.

The recruitment of the HCM cohort, as part of our CIRM cardiomyopathy project, was described elsewhere^24^. Briefly, the targeted patient population were patients with various cardiac procedures and non-cardiac patients with genetic conditions in clinic who were identified to us by their clinical providers. Non-cardiac patients were recruited in person during onsite clinic days or over the phone with permission by the providers. Healthy volunteers were recruited from our cardiovascular prevention clinic (i.e., patients with no diagnosis of heart disease).

### Whole genome sequencing for HCM target cohort

Library preparation and sequencing was performed by Macrogene (first 10 samples) and Novogene on genomic DNA we extracted from iPSC cells (Qiagen DNeasy kit). Paired-end 150bp reads were acquired on the Illumina HiSeq X Ten for a minimum of 90 gigabases of data. Reads were processed using Sentieon’s FASTQ to VCF pipeline (Sentieon version 201808.07)^127^. This pipeline is a drop-in replacement for a BWA^128^ plus GATK best-practices^129^ pipeline for germline SNVs and indels, but has been highly tuned for optimal computational efficiency. BWA alignment to hg38 was followed by deduplication, realignment, base quality score recalibration, and variant calling to generate g.vcf files for each sample. Coverage was assessed (GATK version 3.7)^24^. Individual sample g.vcf files were joined and variant quality score recalibration performed.

### Quality controls of genotype data

We performed stringent quality controls (QCs) for the genotype data following the PRS tutorial (https://choishingwan.github.io/PRS-Tutorial/). For the GWAS summary statistics data (also referred to as the discovery or base data), genetic variants with low minor allele frequency (MAF) and imputation information score (INFO) were removed. We used thresholds suggested in corresponding original papers: MAF < 0.0001, 0.001, 0.0001 and INFO < 0.4, 0.6, 0.6 for T2D, HCM, and AD, respectively. We also excluded duplicated and ambiguous variants to guarantee the accuracy of PRS calculation.

For the individual-level genotype data (also referred to as the target data), we carried out both variant-level and individual-level QCs. We first removed variants with INFO < 0.8 (for UKBB-based cohort), missing call rate > 0.01, MAF < 0.01, or Hardy-Weinberg equilibrium (HWE) < 1 × 10^−6^. For variants with mismatching alleles between discovery and target data, we strand-flipped these alleles to their complementary ones. We further excluded individuals with genotyping rate < 0.01 or with extreme heterozygosity rate (i.e., beyond 3 standard deviations from the mean). Individuals with an up to second degree relative (*π* > 0.125) within the cohort were also removed to prevent bias in prediction evaluation. Eventually, there were 2,176 (*n* = 1088 cases, *n* = 1088 controls), 134 (*n* = 81 cases, *n* = 53 controls), and 1,839 (*n* = 919 cases, *n* = 920 controls) individuals passing the above QCs for T2D, HCM, and AD cohorts, respectively.

To characterize the population structure of target cohorts, principal component analysis (PCA) was performed after pruning (window size = 200 variants, sliding step size = 50 variants, LD *r*^2^ threshold = 0.25). The first 10 principal components (PCs) were retained as covariates in the downstream analysis.

### PLINK C+T PRS calculation

The cell-level C+T PRS was computed using PLINK^31^ which was given by

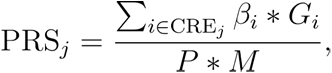

Where CRE*_j_* denotes candidate cis-regulatory elements (CREs) within cell *j*, *β_i_* is the effect size of variant *i*, *G_i_* represents the number of effect alleles, *P* is the ploidy of the sample (2 for human), and *M* is the number of non-missing variants. In the clumping phase, all index variants were forced to be drawn from the variants located within single-cell ATAC-seq peaks of individual cells using the “--clump-index-first” option. Variants within 250 kb of the index variant and three LD thresholds (*r*^2^ = 0.1, 0.3, 0.5) were considered for clumping. After constructing the index variant set, we applied multiple *P*-value thresholds (*P* = 1 × 10^−5^, 1 × 10^−4^, 1 × 10^−3^, 0.01, 0.05, 0.1, 0.5) to compute PRSs, resulting in 21 PRSs calculated for each cell and each individual. We used the 1000 Genomes Project samples to estimate the LD (out-sample estimation) for the simulation and HCM cohorts due to their limited sample sizes, while using the target data (in-sample estimation) for other cohorts.

The standard C+T PRS was calculated using the same set of parameters as that used in computing cell-level PRS, except all variants were considered without conditioning. The *P*-value and LD *r*^2^ thresholds were regarded as hyperparameters in model selection. Similarly, the nonpeak PRS was constructed by limiting the index variants within the genomic regions outside single-cell ATAC-seq peaks.

### Model architecture of scPRS

The cell-level PRS matrix 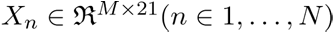 presents single-cell-resolved genetic risk features for each individual, and it is input into the scPRS model to predict the disease risk. Here, *N* and *M* denote the number of individuals and cells, respectively.

scPRS consists of three modules (**Fig. 1**): the feature embedding module, the graph convolutional network module, and the readout module. The feature embedding module takes normalized cell-level PRS *X_n_* as the input and utilizes one-layer perceptron to reweight and integrate 21 PRS features per cell, namely

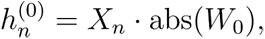

Where *W_O_* is learnable model parameters, abs represents the absolute function, and 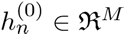 stands for the integrated features of *M* cells for individual *n*. According to the definition of PRS, larger values in *X_n_* indicate higher disease risk. To maintain this interpretability throughout the modeling, we adopt the absolute function abs to enforce nonnegativity for *W_O_*.

We next seek to integrate PRS features across different cells to generate a final risk score. With the consideration of the dropout event and sparsity of single-cell ATAC-seq data, and assuming that cells with similar low-dimensional embeddings should have comparable epigenomes and then similar genetic signals, we employ a graph neural network (GNN)^130^ to smooth and denoise single-cell-level PRS features. More specifically, based on the pre-computed M-kNN graph *G*, the GNN module is defined as

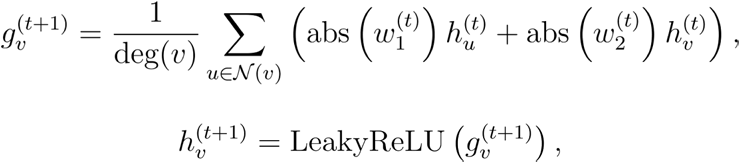

Where 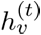 denotes the hidden feature of cell *v* at layer 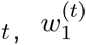 and 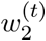 are learnable parameters of layer *t*, ^deg^ denotes the degree of each node/cell, and *N(v)* represents the neighbors of cell *v* in the M-kNN graph *G*. The LeakyReLU activation function is defined as

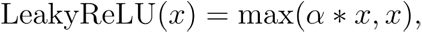

Where *α* = 0.1 is used in this study. Note that the absolute function is also adopted to induce nonnegativity to model weights.

Finally, we design a readout module to map GNN-smoothed hidden features to the phenotype leveraging a one-layer perceptron,

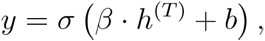

Where 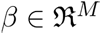 stands for the learnable regression coefficients indicating cell importance to prediction, *T* is the number of total layers in GNN, *b* is the bias term, and *σ* represents the sigmoid function for binary classification and the identify function for regression.

For scPRS+ (cell-level PRSs and non-peak PRSs) and scPRS+covar (cell-level PRSs, non-peak PRSs, age, sex, and 10 PCs), we concatenate additional features to latent cell features *h^(T)^* at the last GNN layer.

### Optimization of scPRS

To train scPRS for disease prediction, we adopt the binary cross-entropy (BCE) loss and additional regularization functions for enhancing prediction power and model interpretability. The loss function ℒ of scPRS is defined as

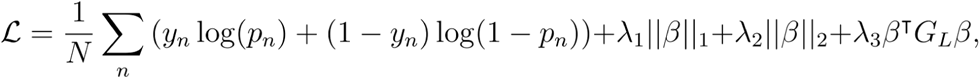

Where 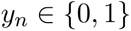 is the true disease label for individual 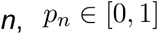 is scPRS-predicted disease probability, 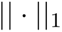 and 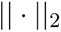 represent *L*_1_ and *L*_2_ norms, respectively. We also add a Laplacian regularization term based on the symmetric normalized Laplacian matrix *G_L_* which is defined as

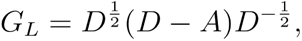

Where D and A denote the degree and adjacency matrices of the cell-cell similarity graph G . We use hyperparameters λ_1_, λ_2_, and λ_3_ to balance across different regularization terms.

scPRS was trained by minimizing the loss ℒ using the Adam algorithm^131^ with learning rate of 1×10^−3^ and batch size of 32. We trained scPRS for 200 epochs. Multiple sets of hyperparameters were considered in model selection, including 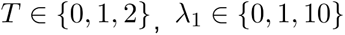, 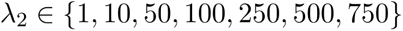, 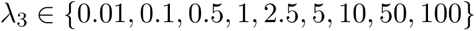, and M-kNN neighbor number 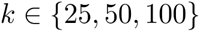 . We also selected between CCA- and Harmony-based cell-cell similarity networks for T2D and AD.

In prediction evaluation, we randomly partitioned the dataset into training, validation, and testing sets comprising 60%, 20%, and 20% of samples, respectively. We trained different scPRS models with all possible combinations of hyperparameters and assessed their performance (measured by AUROC) on the validation dataset. We selected the model yielding the best performance on the validation set and reported its performance on the held-out test set. This process was repeated 10 times to assess the robustness of the model. Prediction performance was evaluated using both the area under the receiver operating characteristic curve (AUROC) and the area under the precision-recall curve (AUPRC).

In cell prioritization, we conducted five-fold cross-validation (CV) and repeated this process five times. The best hyperparameter set was then selected based on the average AUROC score. The final model was trained with this optimal hyperparameter set on the entire dataset. To examine the variability of cell weights learned from model training, we trained 100 models using different random seeds.

For the regression task, the mean squared error (MSE) was used in the loss function instead of BCE. The model performance was evaluated based on the Pearson correlation between true and predicted values.

### LDpred2 and Lassosum

We implemented LDpred2 and Lassosum following the bigsnpr tutorial (https://privefl.github.io/bigsnpr/articles/LDpred2.html). Three LDpred2 models were implemented, including the infinitesimal model (LDpred2-inf), grid model (LDpred-inf), and auto model (LDpred2-auto). All model hyperparameters were selected based on the recommendations provided in the tutorial. To ensure a fair comparison, we maintained the same dataset splits (i.e., train, validation, and test sets) as those used in scPRS. For PLINK C+T, LDpred2-grid, and Lassosum, the best model hyperparameters were determined based on prediction performance on the validation dataset.

### Prioritization of disease-relevant cells and cell types using scPRS

The mapping from input PRS features *X* to latent cell features *h^(T)^* is monotonically increasing, a result of the design principle of scPRS, where weights in the embedding and GNN modules are constrained to be nonnegative. This features facilitates model interpretation: a larger value of *β_m_* denotes a higher enrichment of genetic risk within that cell, thereby informing disease-cell relevance. To account for the variability of learned cell weights, we trained 100 scPRS models and compared the distribution of *β_m_* for individual cells with that of top-ranking weights (i.e., the top 15% of all cell weights per repeat) using one-sided *t*-test. This comparison was conducted for each cell in the dataset. We define “disease-relevant cells” as those cells whose adjusted *P*-values (using the Benjamini-Yekutieli procedure) are less than 0.1. Roughly speaking, scPRS prioritizes cells whose weights are consistently larger than those of the majority of cells.

To get more biological insights, we examined the enrichment of scPRS-prioritized cells within each cell type using Fisher’s exact test. The “disease-relevant cell types” are defined as those cell types whose adjusted *P*-values (using the Benjamini-Hochberg procedure) are less than 0.1.

### Simulation details

Using the PBMC multiome data downloaded from 10x Genomics, we first conducted the differential accessibility analysis to identify monocyte-specific scATAC-seq peaks. In this study, we defined “monocytes” as the total set of CD14/CD16 monocytes and dendritic cells considering their shared heritability^132^. We identified differentially accessible regions (DARs) within monocytes using the top 1500 marker peaks per cell subtype. Next, leveraging a monocyte count GWAS^19^, we computed PLINK C+T PRS conditioned on the variants located within monocyte DARs for a WGS cohort (*n* = 401)^20^. Raw C+T PRS outputs were further standardized to mean = 0 and variance = 1, yielding the “ground-truth” of monocyte count for this cohort.

To introduce randomness, we added a noise term to the simulated monocyte count following

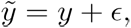

In which 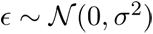. In this study, we used 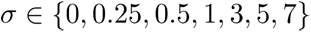. We trained scPRS based on these simulation datasets with and without noises to evaluate its capacity in identifying phenotype-associated cells.

### SCAVENGE

We utilized SCAVENGE^125^ as a benchmark for prioritizing disease-relevant cells. Following the SCAVENGE tutorial (https://sankaranlab.github.io/SCAVENGE/articles/SCAVENGE), we calculated trait relevance scores (TRSs) for individual cells, indicative of their association with the disease. Cells were prioritized by SCAVENGE if their TRSs were above 95% of all TRSs. As in the scPRS analysis, we evaluated the enrichment of selected cells within each cell type using Fisher’s exact test.

### Stratified LDSC

Partitioned heritability analysis was carried out using the stratified LD score regression (sLDSC) as previously described^36^. Heritability was quantified within the total set of snATAC-seq peaks identified for each of the left ventricle cell types. Genetic enrichment for a particular cell type was defined by calculating the captured heritability per unit of sequence within the total set of identified snATAC-seq peaks for that cell type, compared to the genome overall. *P*-values were calculated as previously described^36^; nominal significance (*P* < 0.05) was taken to be indicative of true enrichment.

### Identification of disease-relevant CREs

As the first step of the layered multiomic analysis (**Fig. 5a**), we identified differentially accessible CREs within each scPRS-prioritized cell type using Signac^66^. Specifically, we employed the FindMarker function to compare peaks within scPRS-prioritized cells (per cell type) against the background, with parameters test.use=’LR’, latent.vars=’peak_region_fragments’, min.pct = 0.02, and logfc.threshold=0.1. Significant peaks (adjusted *P* < 0.1 based on BH correction) with a positive log_2_ fold change were defined as differentially accessible CREs. Next, leveraging the discovery GWAS summary statistics, we conducted MAGMA^55^ analysis for these differentially accessible CREs per cell type, with gene-model=’multi’. We defined disease-relevant CREs (T2D-CREs and AD-CREs) as those CREs with adjusted MAGMA *P* < 0.1 based on BH correction. We expanded our analysis to involve all nominally significant CREs (MAGMA *P* < 0.05) for HCM, as no CRE passed the multiple testing correction.

### Mapping CRE-gene links

We mapped CREs to their target genes based on two complementary strategies. First, we adopted the closest-gene strategy^133^ and assigned each CRE to its closest gene. In addition, we added more distant genes based on a coaccessibility analysis using Cicero^56^ and linked each CRE to those genes whose TSS peak displayed coaccessibility with the CRE above 80% of all interactions. For each scPRS-prioritized cell type, the expressed genes mapped to disease-relevant CREs within that cell type define the repertoire of disease candidate genes.

### Enrichment of disease-associated variants within scPRS-cell-specific peaks

Per disease-relevant cell type, we performed clumping within differentially accessible peaks in scPRS-prioritized cells to remove redundant variants. Multiple LD *r*^2^ thresholds (*r*^2^ = 0.1, 0.3, 0.5) were tested. Leveraging the clumped variant set, we examined the enrichment of disease-associated variants (GWAS *P* < 5 × 10^−8^) within scPRS-cell-specific peaks by comparing it with the genome-wide distribution.

### Transcription factor binding motif analysis

The TF binding motif analysis was performed using GimmeMotifs^134^. The differential motifs between disease-relevant CREs and all peaks within the corresponding cell type were identified using the “gimme motif” command with options -f=0.5 and -s=0. AUROC was adopted to quantify the motif enrichment.

### Network analysis

We downloaded the human protein-protein interactions (PPIs) from STRING v12.0^80^, comprising 19,622 proteins and 6,857,702 interactions. High-confidence PPIs (combined score >700) were extracted for downstream analysis, including 16,185 proteins and 236,000 interactions. To mitigate bias from hub proteins^135^, we applied the random walk with restart (RWR) algorithm with a restart probability of 0.5. This produced a smoothed network after retaining the top 5% predicted edges (*n* = 6,243,766). Next, we utilized the Louvain method^136^ to decompose the network into different modules. Following algorithm convergence, we obtained 1,261 modules with an average size of 13 nodes.

The enrichment of genes of interest within each module was tested using the hypergeometric test. Modules with adjusted *P* < 0.1 based on BH correction were considered significant.

### scVINet design and training

scVINet was trained to predict CREs across various cell types based on the DNA sequence (**Extended Data Fig. 4c**). Specifically, scVINet takes a 2000-bp DNA sequence as the input, and outputs the CRE status of the centered 200 bp for different cell types. The CRE label for a specific cell type is 1 if over 50% of the centered 200 bp is overlapped by an ATAC-seq peak within that cell type, and is 0 otherwise. scVINet model structure follows the Beluga architecture^59^, except its outputs correspond to different cell types within the tissue of interest.

CREs within chromosomes 6 and 7, and chromosomes 8 and 9 were held out as validation and test data, respectively. CREs in other chromosomes were used as training data. Genomic regions annotated by the ENCODE blacklist^137^ were excluded from analysis. We adopted the BCE loss as the objective function. scVINet was trained using the stochastic gradient descent (SGD) algorithm with weight decay coefficient of 1 × 10^−6^, momentum of 0.9, learning rate of 0.08, and batch size of 64. scVINet was implemented using Selene^60^, a PyTorch-based library for sequence deep learning modeling. In this study, we developed three different versions of scVINet specified as scVINet-panc, scVINet-heart, and scVINet-brain by training scVINet on the human pancreas, left ventricle, and cortex single-cell ATAC-seq data, respectively.

### Prediction of variant functional effects using scVINet

We employed scVINet to predict the impact of genetic variants on the functionality of CREs across diverse cell types. For a given cell type *c* and variant *v* (from reference allele to alternative allele), scVINet predicts the status of CRE *y*_ref,*c*_ and *y*_alt,*c*_ for sequences centered on the reference and alternative alleles, respectively. We define the functional effect of variant *v* in cell type *c* as *y_v_*_,*c*_ = *y*_alt,*c*_ - *y*_ref,*c*_, representing how the variant alters CRE in this cell type. To achieve a global evaluation of functional scores, we introduce the *Z*-score *Z_v_*_,*c*_ which normalizes *y_v_*_,*c*_ as *Z_v_*_,*c*_ = (*y_v_*_,*c*_ - *μ*) / *σ*, where *μ* and *σ* denote the mean and standard deviation of all variant scores, respectively. The *Q*-score *Q_v_*_,*c*_ is further defined as the quantile of the absolute *Z*-score |*Z_v_*_,*c*_| among all variants. A higher *Q*-score indicates a larger functional effect within a specific cell type.

### Benchmarking scVINet prediction

To benchmark scVINet prediction on variant effects against QTL analysis (eQTL or caQTL), we compared the absolute *Z*-scores computed by scVINet between QTLs and non-QTLs using a two-sided *t*-test. The *t*-statistics was used to measure the enrichment of functional variants defined by scVINet within QTLs.

As the second benchmarking, we employed SNP2TFBS^71^ to predict the effects of variants on altering TF binding site affinity. The binding affinities for different TFs were averaged for each studied variant to estimate its overall effect. Given a particular quantile cutoff, variants were split into two groups according to their *Q*-scores. We then compared the averaged SNP2TFBS scores between these two groups of variants using a two-sided *t*-test. We reported *t*-statistics which indicates the enrichment of TFBS disrupting variants within scVINet-defined functional variants.

### Variant effect within disease-relevant CREs

We compared the abundance of functional disease-associated variants (GWAS *P* < 0.05) within disease-relevant CREs against the background using Fisher’s exact test. Similarly, the functional variants were defined as those with *Q*-scores above a given cutoff (multiple cutoffs applied). Odds ratio was adopted to measure the enrichment of functional variants within disease-CREs.

### Fine-mapping disease risk variants

We utilized three approaches to fine-map disease risk variants, including scVINet, QTL, and TFBS. A 0.8 quantile cutoff was adopted to define functional variants based on scVINet in fine-mapping. In addition to SNP2TFBS, motifbreakR^73^ was also used to predict variant disruption on TF binding. A positive averaged SNP2TFBS score or a strong-effect motifbreakR score was used to define a disrupting variant. We excluded missense and loss-of-function (LoF) variants, and variants with GWAS *P* >= 0.05 from fine-mapping.

### iPSC reprogramming

Induced pluripotent stem cells (iPSCs) were reprogrammed from PBMCs using Sendai virus (CytoTune iPS 2.0 Sendai Reprogramming Kit) as previously described^138^. Three clones were generated per subject, karyotyped (KaryoStat, ThermoFisher Scientific), determined to be mycoplasma-free, and evaluated by immunohistochemistry for expression of pluripotency markers TRA-1-60 (LifeTech MA1023) and SSEA4 (LifeTech MA1021). Cells were maintained under feed-free conditions in mTeSR (STEMCELL Technologies, 5850) or Essential 8 media (Fisher, A1517001) and stored in liquid nitrogen.

### Cardiomyocyte differentiation and drug treatment

As previously described^139^, iPSCs were plated on Matrigel and cultured in StemMACS iPS-Brew XF (MACS Miltenyi Biotec, 130-104-368) until the final passage in Essential 8 media (Fisher, A1517001). Cardiomyocyte differentiation was induced at 60-80% confluency, with culture in RPMI media (Gibco/LifeTech 11875-119) plus B27 supplement lacking insulin (Gibco/LifeTech A1895601). 6µM of CHIR-99021 (Fisher, NC0976209) was added on day 0 and 6 µM IWR1 (Fisher, NC1319406) was added on day 3. Beginning on day 7, media was changed every other day using RPMI media supplemented with B27 containing insulin (Gibco/LifeTech 17504-044). Upon commencement of beating (around day 15), cells underwent purification via a three-day glucose starvation (RPMI media without glucose [Gibco/LifeTech 11879-020] supplemented with insulin-containing B27), a one-day recovery in glucose-containing media, and subsequent replating (dissociated in TrypLE, Fisher, 50-591-353). Cells were then maintained in RPMI media supplemented with insulin-containing B27 until approximately day 30. After differentiation, drug treatment occurred at 0 hours and 24 hours and samples assayed at 48 hours. Cells were treated with 250nM MYK-461 (Cayman Chemical, 19216-5mg), 400nM or 1uM omecamtiv mecarbil (Selleckchem via Fisher, NC1069600), or DMSO.

### RNA-seq library preparation, sequencing, quality control, and expression matrix generation

RNA was extracted from iPSCs or cardiomyocytes (RNeasy, Qiagen). Illumina RNA-seq libraries (TruSeq Stranded Total RNA LP Gold) were prepared on the Bravo (Agilent), pooled, and sequenced (NovaSeq-6000, paired-end, 100bp)^24^. Where possible drug treatment conditions for the same differentiation were kept together in batches, while replicate differentiations for the same iPSC lines were split apart, and HCM and control samples were distributed across batches. Reads were aligned to hg38 (STAR). Principal component analysis on cardiomyocyte and iPS samples separately returned no outlier samples (as defined as *Z*-score of principal component 1 > 3). Library quality control was assessed via fastp, fastQC, STAR, and Picard metrics. Samples were flagged for poor QC by the following metrics: GC content after filtering outside of 20-80% (fastp), duplication rate greater than 40% (fastp), uniquely mapped read pairs (fragments) < 20 million (STAR), mean reads (average of forward and reverse) < 20 million (fastQC), ribosomal RNA bases > 20% (Picard), coding plus UTR (untranslated region) < 50% (Picard), uniquely mapping fragments < 60% (STAR). Samples with more than one flag were removed. Cardiomyocyte and iPSC samples were subsequently processed separately. Reads were computed as CPM (edgeR) and corrected for library preparation batch (combat-seq) and TMM normalized (edgeR) to generate the final expression matrix. For samples with biological replicates, TMM counts were averaged. Principal component analysis was performed and principal component 1 assessed for spearman correlation with the following metadata: percent GC content (fastp), mean reads (average of forward and reverse) in millions (fastQC), percent ribosomal RNA bases (Picard), uniquely mapped fragments in millions (STAR), duplication rate (fastp), percent coding or UTR (picard), library preparation batch, and sequencing pool. The maximum absolute value for spearman correlation between PC1 and the library metadata was 0.08 for cardiomyocyte samples, indicating good quality control with technical artifacts having minimal influence on the dataset. iPSC samples had higher correlation for three metrics (0.26 with GC content, 0.22 with duplication rate, and 0.11 with percent coding or UTR), with the remaining less than an absolute value of 0.04.

### Differential expression analysis

Raw data was input into DESeq2^140^ as required to compare gene expression between HCM cases and controls across different conditions. Gene counts were averaged across replicates. Sample sex and ancestry were included as covariates in the analysis.

### Allelic imbalance analysis in rs7922621 prime-edited microglia

rs7922621 prime-edited WTC11 clones were obtained from our previous study^90^ and microglia were differentiated accordingly. Total RNA was isolated from wild-type and prime-edited microglia using the RNeasy plus mini kit (Qiagen, 74034). 400 ng of total RNA was reverse transcribed using the iScript cDNA Synthesis kit (Bio-Rad, 1708891). The cDNA region containing phased heterozygous SNP of *ANXA11* – rs2573353 in WTC11^141^ – was amplified using the following primers: WTC-ANX-F: AGGTCCAATAATCCCTGCTGA, WTC-ANX-R: CCATGGTGCTCGGCTAATTT. The PCR products were purified by agarose gel extraction, followed by addition of Illumina adaptors and deep sequencing. Reads were aligned to the sequence of either allele and counted if the 100 bp region surrounding rs2573353 were exactly matched.

### Differentiation of TMEM119-Tdtomato reporter cell line induced-microglia

iPSCs stably expressing a TMEM119-tdTomato reporter transgene were first differentiated into fibroblast-like cells using a previously established method^92,93^. TMEM119-tdTomato fibroblasts were seeded onto 96-well plates (Corning) coated with 0.1% gelatin and Matrigel in fibroblast medium (DMEM with 10% FBS and 1% penicillin/streptomycin). After 48 hours, the cells were transduced with 200μL of two different concentrated retroviruses to overexpress the human *PU.1* and *CEBPA* per 96-well well with 5μg/ml polybrene in fibroblast medium. Twenty-four hours after transduction, media was switched to DMEM with 5% FBS, human M-CSF 10 ng/ml and IL-34 10 ng/ml, and media was refreshed every three days thereafter. Induced microglia-like cells (iMGs) expressing the TMEM119-tdTomato reporter were used for experiments 14 days after viral transduction.

### siRNA transfection

siRNAs (Thermofisher) at a concentration of 30nM were transfected into iMGs on day 14 using Lipofectamine™ RNAiMAX Transfection Reagent (Thermofisher Scientific, 13778075) in complete iMG medium (DMEM +5% FBS, 10ng/mL M-CSF, and IL-34 10ng/mL). After 24 hours, the medium was refreshed with complete iMG medium, and after an additional 24 hours (48 hours post transfection), cell cultures were collected for qRT-PCR or pHrodo analysis.

### pHrodo phagocytosis assay

iMGs cultured in 96-well plates (Corning) coated with Matrigel and gelatin were incubated with 10μg of pHrodo Green *E. coli* Bioparticles (Inucyte) for 15 minutes at 37°C. Wells were then washed with PBS and were longitudinally imaged with Molecular Devices ImageExpress at 30-minute intervals for the initial 2 hours and 1-hour intervals up to 24 hours after the start. The 2-hour time point was selected for downstream analysis. ImageJ software was used for quantification of individual replicates across conditions based on the co-localization of the TMEM119-Tdtomato and pHrodo green.

## Extended Data

**Extended Data Fig. 1.**
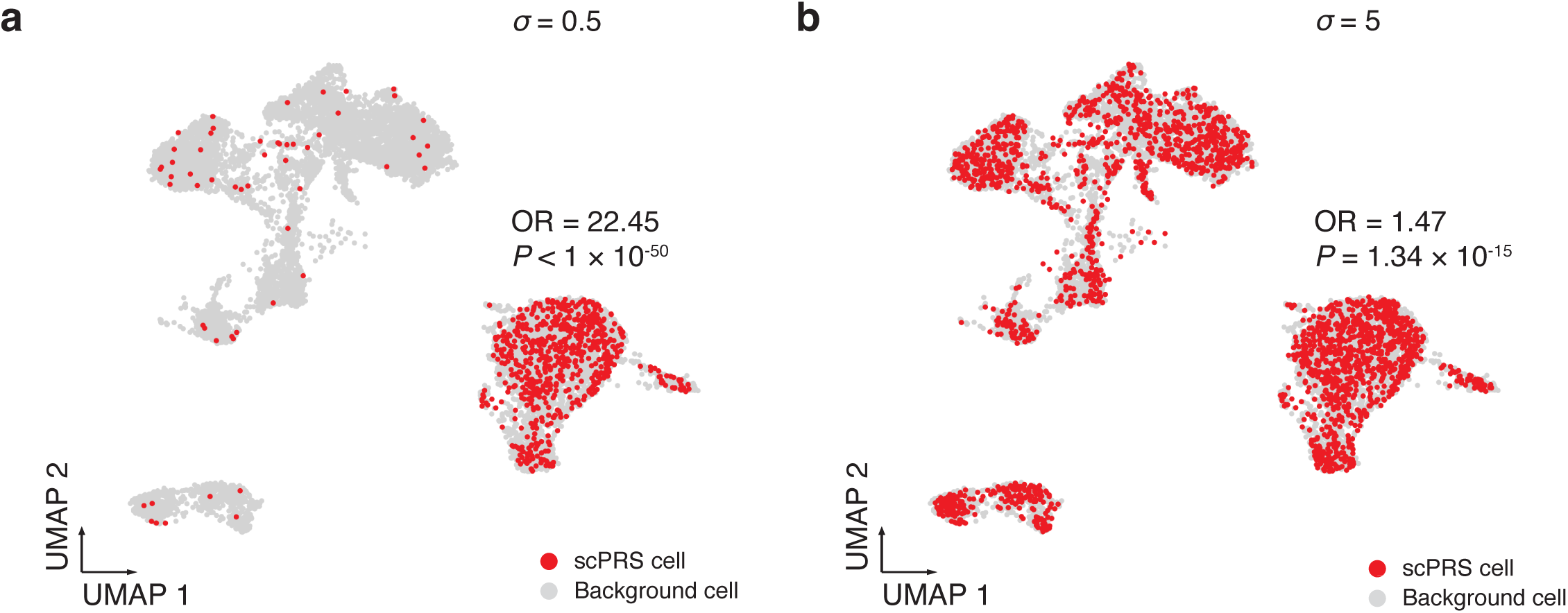
Additional simulation results. **a-b**, Monocyte-count-relevant cells prioritized by scPRS (in red) in different noise settings: *σ* = 0.5 (**a**) and *σ* = 5 (**b**). *P*-value and *Z*-score by two-sided Fisher’s exact test. UMAP, uniform manifold approximation and projection; OR, odds ratio.

**Extended Data Fig. 2.**
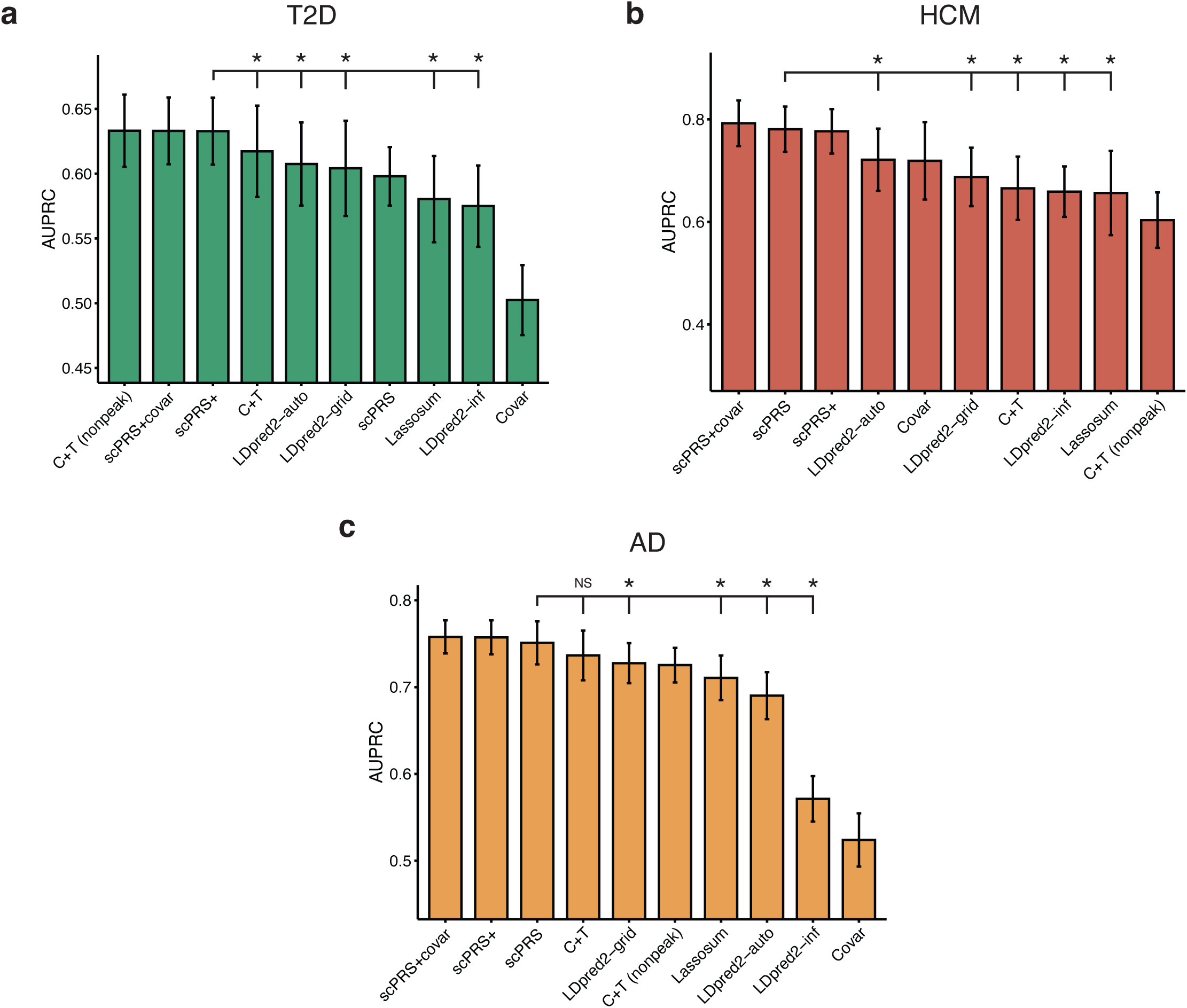
Benchmark results on disease prediction. **a-c**, Barplots of AUPRC scores (*n* = 10 repeats) of different models for T2D (**a**), HCM (**b**), and AD (**c**), respectively. Training, validation, and test data splits were kept identical across different methods to enable rigorous comparison. scPRS+, scPRS model integrating non-peak PRSs; scPRS+covar, scPRS model integrating non-peak PRSs and covariates; C+T, clumping and thresholding PRS; C+T (nonpeak), logistic regression model of non-peak C+T PRSs; Covar, logistic regression model of covariates. Performance comparison was conducted using one-sided paired *t*-test. *, adjusted *P* < 0.1; NS, not significant; AUPRC, the area under the precision-recall curve. The mean and 95% confidence interval (CI) are annotated using the barplot and error bar, respectively.

**Extended Data Fig. 3.**
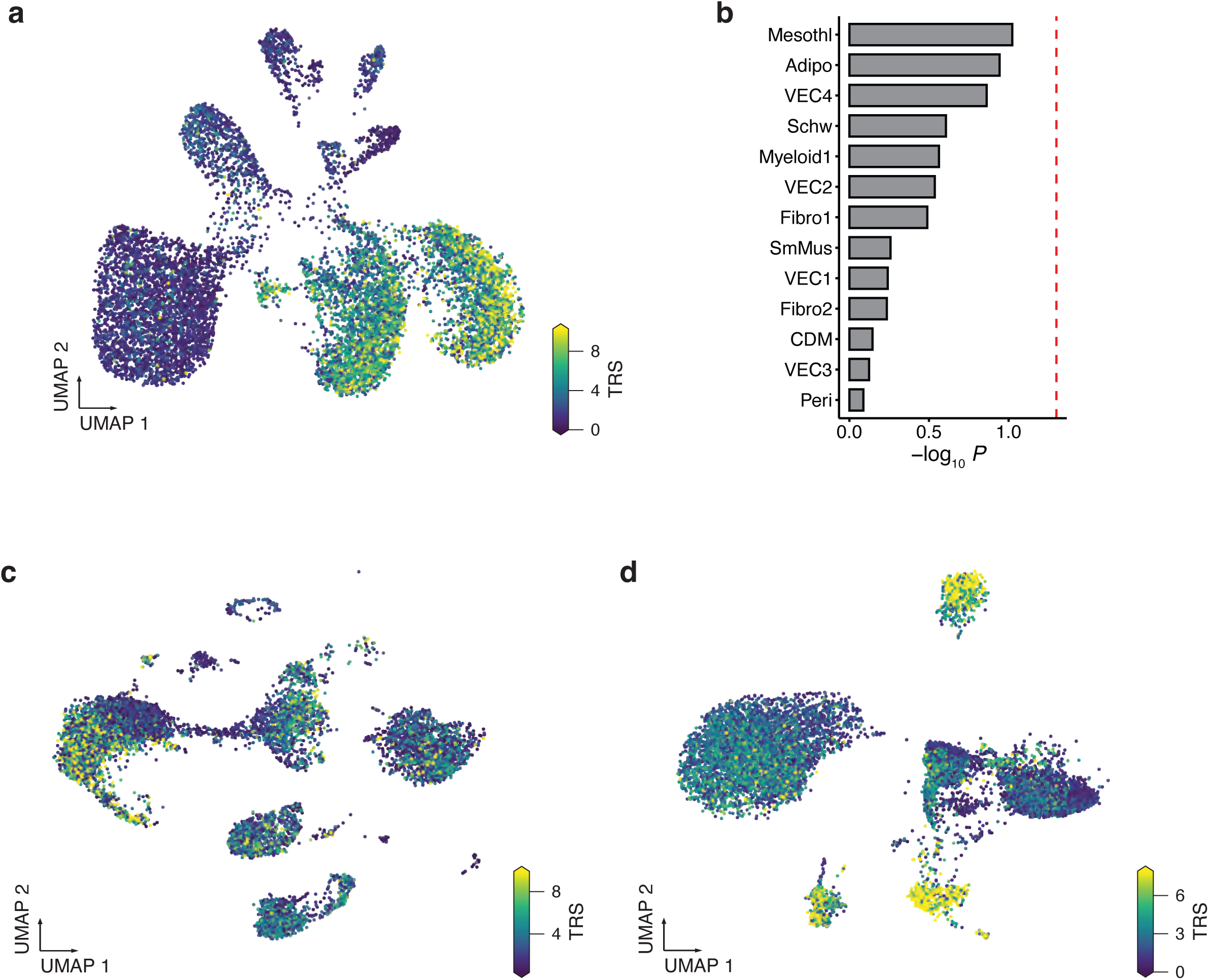
Benchmark results on disease cell prioritization. **a**, T2D-cell relevance computed by SCAVENGE. UMAP, uniform manifold approximation and projection; TRS, trait relevance score. **b**, The enrichment of HCM heritability within different cell types. The red dashed line indicates *P* = 0.05. *P*-value by stratified LDSC (sLDSC). Mesothl, mesothelial cell; Adipo, adipose cell; CDM, cardiomyocyte; Fibro, fibroblast; LEC, lymphatic endothelial cell; Peri, pericyte; Schw, Schwann cell; SmMus, smooth muscle cell; VEC, vascular endothelial cell. **c-d**, Disease-cell relevance computed by SCAVENGE for HCM (**c**) and AD (**d**), respectively. The same GWAS and single-cell datasets were used in SCAVENGE as those used in scPRS to ensure comparison fairness.

**Extended Data Fig. 4.**
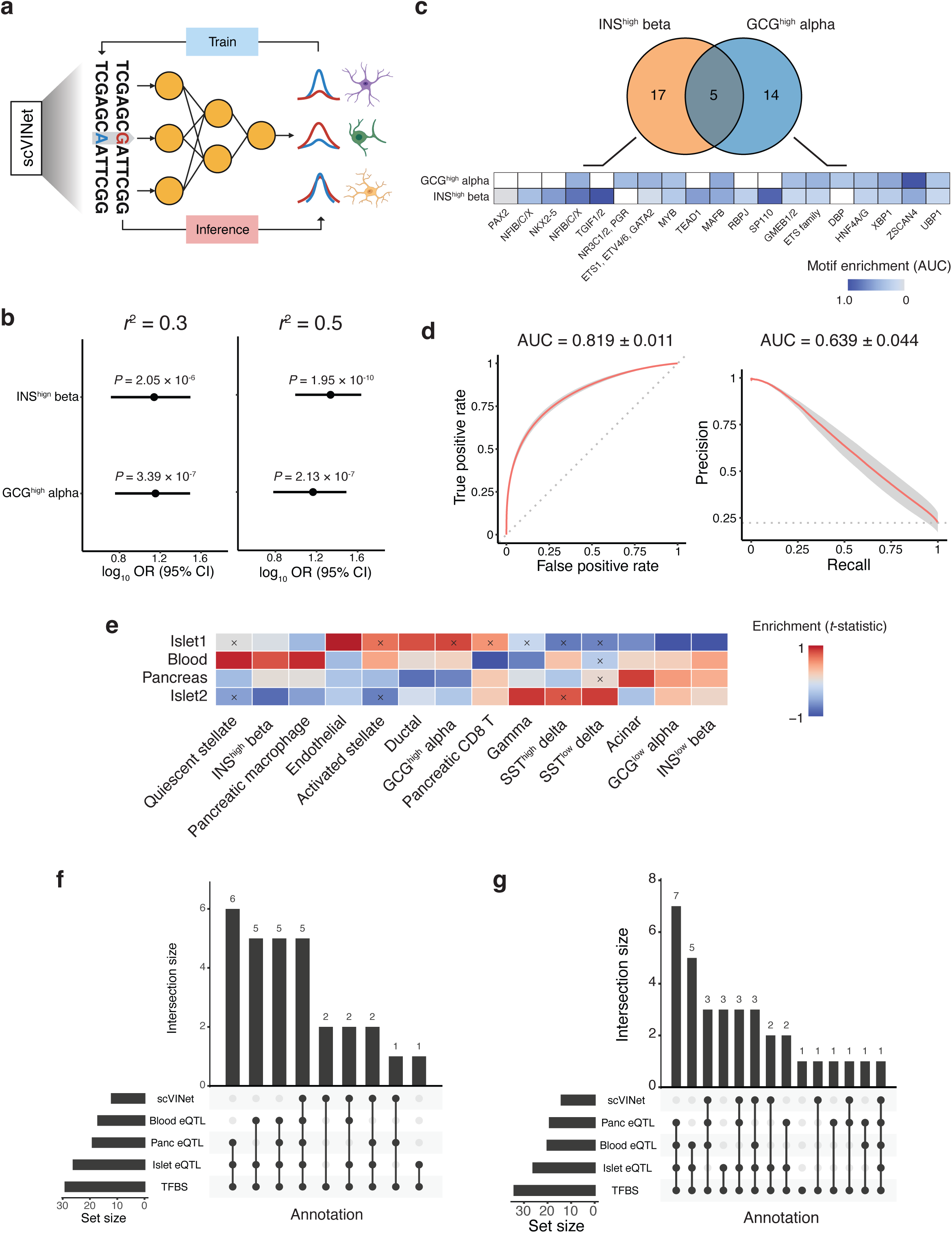
Additional results of multiomic analysis for T2D. **a**, Schematic of the scVINet model. **b**, Enrichment of T2D-associated variants within CREs differentially accessible in scPRS-prioritized cells. Two linkage disequilibrium (LD) thresholds, *r*^2^ = 0.3 (left) and *r*^2^ = 0.5 (right), were adopted in clumping. *P*-value by two-sided Fisher’s exact test. The log_10_(OR) and 95% CI are annotated by the dot and error bar, respectively. OR, odds ratio; CI, confidence interval; CRE, cis-regulatory element. **c**, T2D-CREs (top) and their motif enrichment (bottom) in two T2D-relevant cell types including GCG^high^ alpha and INS^high^ beta cells. Motif enrichment was measured by AUC. Row-wise standardization was performed. Only significant enrichment (adjusted *P* < 0.1, Bonferroni correction) is colored. AUC, the area under the receiver operating characteristic (ROC) curve. **d**, The receiver operating characteristic curve (ROC; left) and the precision-recall curve (PRC; right) showing scVINet-panc test performance across various cell types in the human pancreas. The red line and gray area represent the mean and 95% CI, respectively. The dashed gray line indicates the random prediction. AUC, the area under the curve. **e**, Comparison of scVINet-panc prediction scores between eQTLs and other variants across different tissues and cell types. Comparison was performed using two-sided *t*-test. ×, adjusted *P* > 0.1 by Benjamini-Hochberg correction. Columns were standardized. **f-g**, Summary statistics of prioritized T2D risk variants in GCG^high^ alpha cells (**f**) and INS^high^ beta cells (**g**) using different annotations. TFBS, transcription factor binding site; Panc, pancreas.

**Extended Data Fig. 5.**
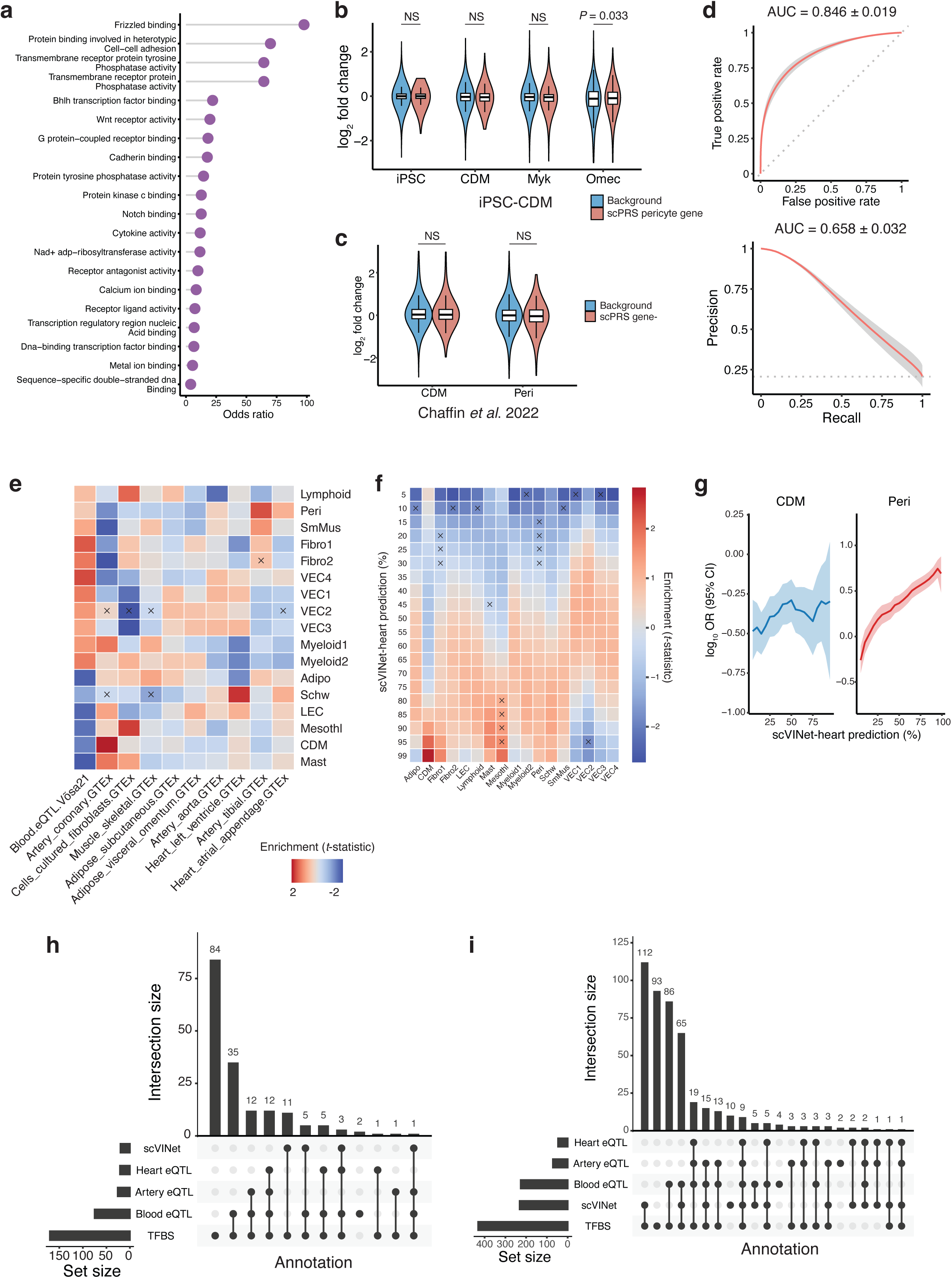
Additional results of multiomic analysis for HCM. **a**, Lollipop chart of gene ontology (GO) enrichment (molecular function) for M16 genes. Significant GO terms (adjusted *P* < 0.1 by Benjamini-Hochberg (BH) correction) are shown. **b**, Expression fold change comparison between HCM pericyte genes and background transcriptome based on the HCM iPSC RNA-seq data. *P*-value by two-sided *t*-test. The box plot center line, limits, and whiskers represent the median, quartiles, and 1.5x interquartile range (IQR), respectively. CDM, cardiomyocyte; Myk, Mavacamten; Omec, Omecamtiv mecarbil; iPSC, induced pluripotent stem cell; NS, not significant. **c**, Expression analysis of HCM cardiomyocyte and pericyte genes based on the pericyte and cardiomyocyte transcriptome data, respectively. Peri, pericyte. **d**, The receiver operating characteristic curve (ROC; top) and the precision-recall curve (PRC; bottom) showing scVINet-heart test performance across various cell types in the human left ventricle. The red line and gray area represent the mean and 95% CI, respectively. The dashed gray line indicates the random prediction. AUC, the area under the curve; CI, confidence interval. **e**, Comparison of scVINet-heart prediction scores between eQTLs and other variants across different tissues and cell types. Comparison was performed using two-sided *t*-test. ×, adjusted *P* > 0.1 by BH correction. Rows were standardized. Mesothl, mesothelial cell; Adipo, adipose cell; CDM, cardiomyocyte; Fibro, fibroblast; LEC, lymphatic endothelial cell; Peri, pericyte; Schw, Schwann cell; SmMus, smooth muscle cell; VEC, vascular endothelial cell. **f**, Enrichment of transcription factor binding site (TFBS) disrupting variants within scVINet-heart-prioritized variants (various thresholds applied). Enrichment was quantified by *t*-statistics. ×, adjusted *P* > 0.1 by BH correction. **g**, Enrichment of scVINet-heart-prioritized HCM-associated variants within HCM-CREs. OR, odds ratio. OR and CI by two-sided Fisher’s exact test. OR is annotated by the solid line and 95% CI is represented by the shaded area. **h-i**, Summary statistics of prioritized HCM risk variants in cardiomyocytes (**h**) and pericytes (**i**) using different annotations.

**Extended Data Fig. 6.**
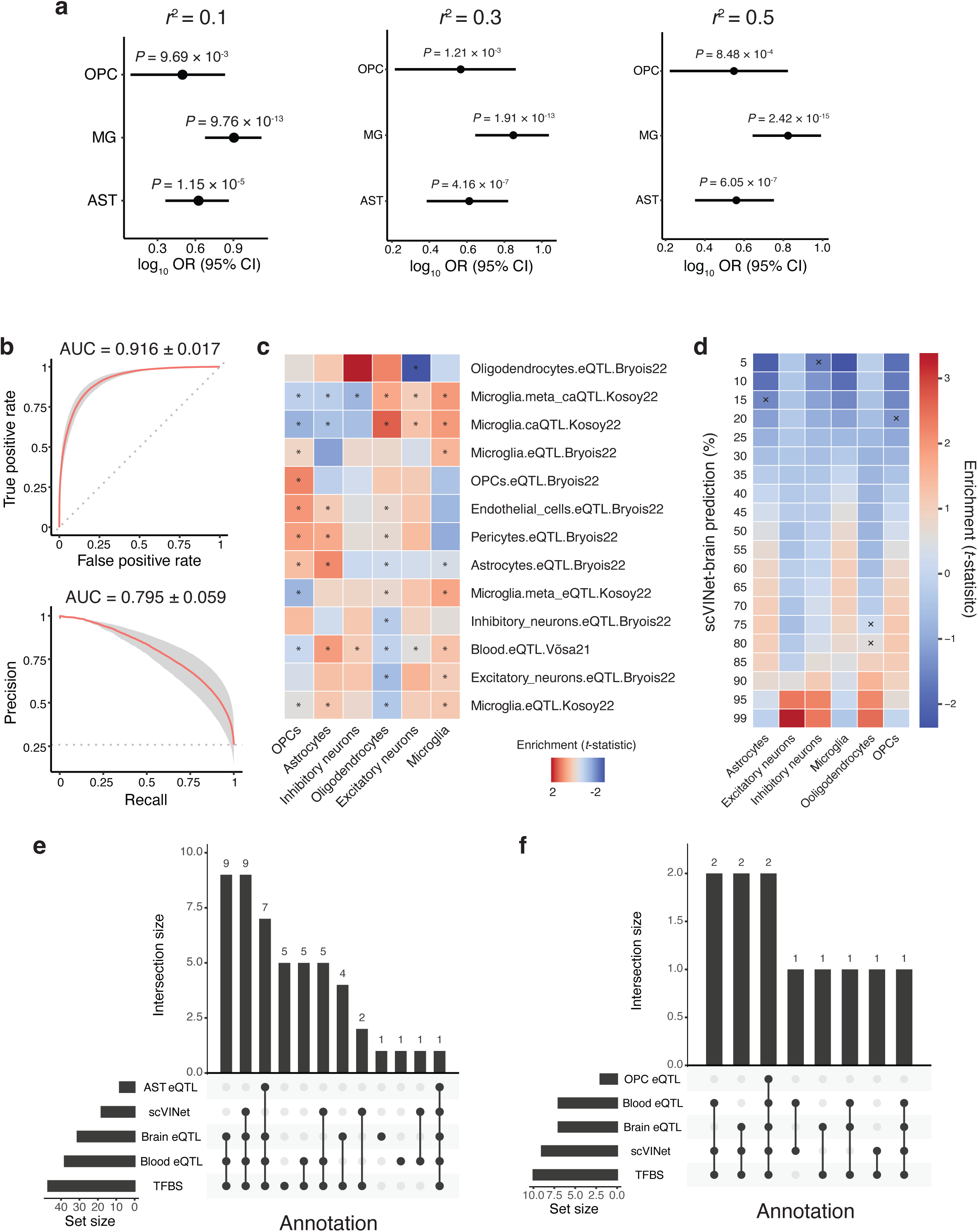
Additional results of multiomic analysis for AD. **a**, Enrichment of AD-associated variants within CREs differentially accessible in scPRS-prioritized cells. Different linkage disequilibrium (LD) thresholds, including *r*^2^ = 0.1 (left), *r*^2^ = 0.3 (middle), and *r*^2^ = 0.5 (right), were adopted in clumping. *P*-value by two-sided Fisher’s exact test. The log_10_(OR) and 95% CI are annotated by the dot and error bar, respectively. AST, astrocyte; MG, microglia; OPC, oligodendrocyte progenitor cell; OR, odds ratio; CI, confidence interval; CRE, cis-regulatory element. **b**, The receiver operating characteristic curve (ROC; top) and the precision-recall curve (PRC; bottom) showing scVINet-brain test performance across various cell types in the human cortex. The red line and gray area represent the mean and 95% CI, respectively. The dashed gray line indicates the random prediction. AUC, the area under the curve. **c**, Comparison of scVINet-brain prediction scores between eQTLs and other variants across different tissues and cell types. Comparison was performed using two-sided *t*-test. *, adjusted *P* < 0.1 by Benjamini-Hochberg (BH) correction. Rows were standardized. **d**, Enrichment of transcription factor binding site (TFBS) disrupting variants within scVINet-brain-prioritized variants (various thresholds applied). Enrichment was quantified by *t*-statistics. ×, adjusted *P* > 0.1 by BH correction. **e-f**, Summary statistics of prioritized AD risk variants in astrocytes (**e**) and OPCs (**f**) using different annotations.

